# Matrix factorization recovers consistent regulatory signals from disparate datasets

**DOI:** 10.1101/2020.04.26.061978

**Authors:** Anand V. Sastry, Alyssa Hu, David Heckmann, Saugat Poudel, Erol Kavvas, Bernhard O. Palsson

## Abstract

The availability of gene expression data has dramatically increased in recent years. This data deluge could result in detailed inference of underlying regulatory networks, but the diversity of experimental platforms and protocols introduces critical biases that could hinder scalable analysis of existing data. Here, we show that the underlying structure of the *E. coli* transcriptome, as determined by Independent Component Analysis (ICA), is conserved across multiple independent datasets, including both RNA-seq and microarray datasets. We also show that echoes of this structure remain in the proteome, accelerating biological discovery through multi-omics analysis. We subsequently combined five transcriptomics datasets into a large compendium containing over 800 expression profiles and discovered that its underlying ICA-based structure was still comparable to that of the individual datasets. ICA thus enables deep analysis of disparate data to uncover new insights that were not visible in the individual datasets.

## Introduction

Publicly available datasets, such as the NCBI Gene Expression Omnibus (GEO)^1^ and Array Express^2^, contain thousands of transcriptomics datasets that are often designed and analyzed for a specific study. Historically, microarrays were the platform of choice for transcriptomic interrogation. Over the past decade, usage of RNA sequencing (RNA-seq) has surpassed microarrays due to its higher sensitivity and ability to detect new transcripts^3^. Public repositories for proteomics datasets have introduced additional reusable expression datasets^4,5^.

Multiple consortia have performed extensive comparisons of expression levels across different microarray and RNA-seq platforms^6–8^. These studies showed that absolute gene expression levels cannot be accurately measured by either expression profiling technique, whereas relative abundances are consistent across a wide range of transcriptomics platforms, with appropriate quality controls. However, transcript levels alone cannot predict protein expression levels^9^To further complicate matters, batch effects and technical heterogeneity continue to present significant challenges to successful integration of omics datasets^10^.

Differential expression analysis is the most common analytical method applied to transcriptomics datasets. However, differential expression analysis is limited in dimensionality, interpretability, and reproducibility; it can only be applied to pairs of experimental conditions, requires additional analysis to interpret large swaths of differentially expressed genes^11,12^, and is highly dependent on the quantification pipeline^13,14^. Alternatively, machine learning methods, especially matrix factorization^15,16^, have provided new tools for extracting low-dimensional biological information from large omics data.

In particular, independent component analysis (ICA) has been shown to extract biologically significant gene sets from many transcriptomics datasets^17–22^ and two proteomics datasets^23^. ICA outperformed 42 module detection methods in a comprehensive examination across 5 organisms^24^. Previously, we applied ICA to a high-quality *Escherichia coli* gene expression compendium to extract 92 independently-modulated groups of genes (called i-modulons). Of these 92 i-modulons, 61 represented transcriptional regulators and their compendium-wide activities. An additional 25 i-modulons were linked to biological functions or genetic perturbations, leaving only 6 uncharacterized. I-modulons have provided clear biological explanations for transcriptional changes in significantly perturbed cells^25,26^. Other research groups have successfully identified co-regulated gene sets using ICA on human transcriptomic datasets^19,22^, but characterization of some components was hindered due to the high fraction of unknown human genes compared to the model bacterium *E. coli*^27,28^.

A recent study showed that among dimensionality reduction methods, ICA produced the most similar components across 14 independent cancer microarray datasets^29^. In this study we show that consistent regulatory components can be identified in expression datasets spanning disparate experimental conditions, and that these components are robust to dataset integration. Through analysis of five independent *E. coli* transcriptomics datasets and two independent proteomics datasets, we identify a coherent structure without requiring batch normalization procedures. In addition, integrated analysis of the different datasets demonstrated compelling evidence towards regulon discovery. These results present ICA as a promising tool to integrate and understand the flood of omics data challenging scientists today.

## Results

### ICA identifies meaningful “i-modulons” in five transcriptomic datasets

We compiled two RNA-seq and three microarray datasets, each using a different expression profiling technology or generated from a different research group (Table 1, Supplementary Data 1). Each dataset was independently processed, centered, and decomposed with ICA (See Methods). This process generated a set of independent components (ICs) for each dataset (Figure 1a, Supplementary Data 2). Each IC contains a coefficient for every gene, although most gene coefficients were near zero for a given IC. To understand the biological role of each IC, we selected genes with sufficiently non-zero coefficients and refer to this set of genes as an “i-modulon”^30^. ICs, and their corresponding i-modulons, represent the underlying structure of the transcriptome under any experimental condition in the database. The condition-dependent dynamics of gene expression are captured by the activities of the ICs (also referred to as i-modulon activities, Supplementary Data 3). In this study, we focused on the condition-invariant structure of the transcriptome (i.e. ICs and their resultant i-modulons), to determine if transcriptome structure is conserved across platforms.

**Table 1:**
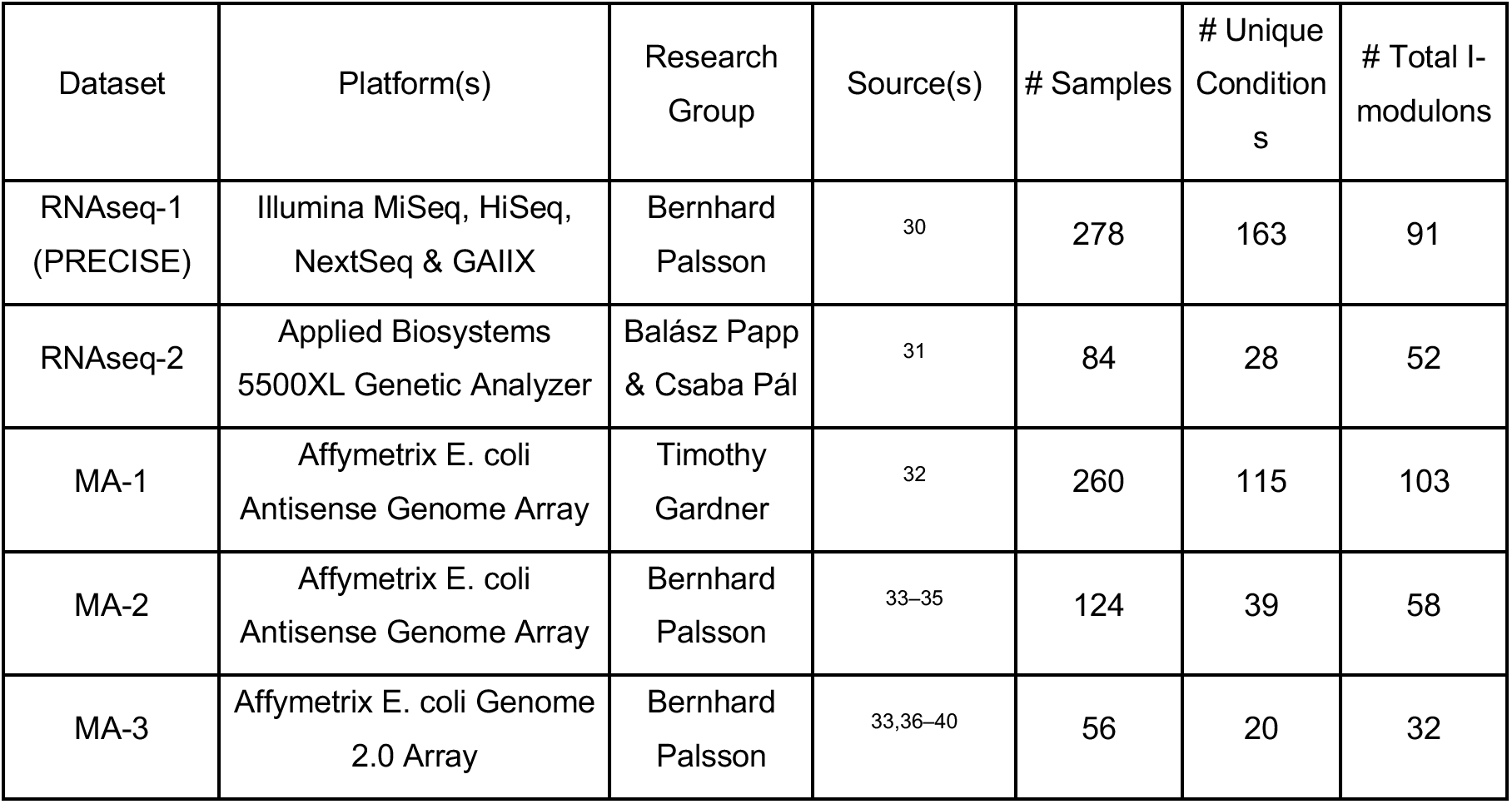
Summary of transcriptomic datasets

**Figure 1:**
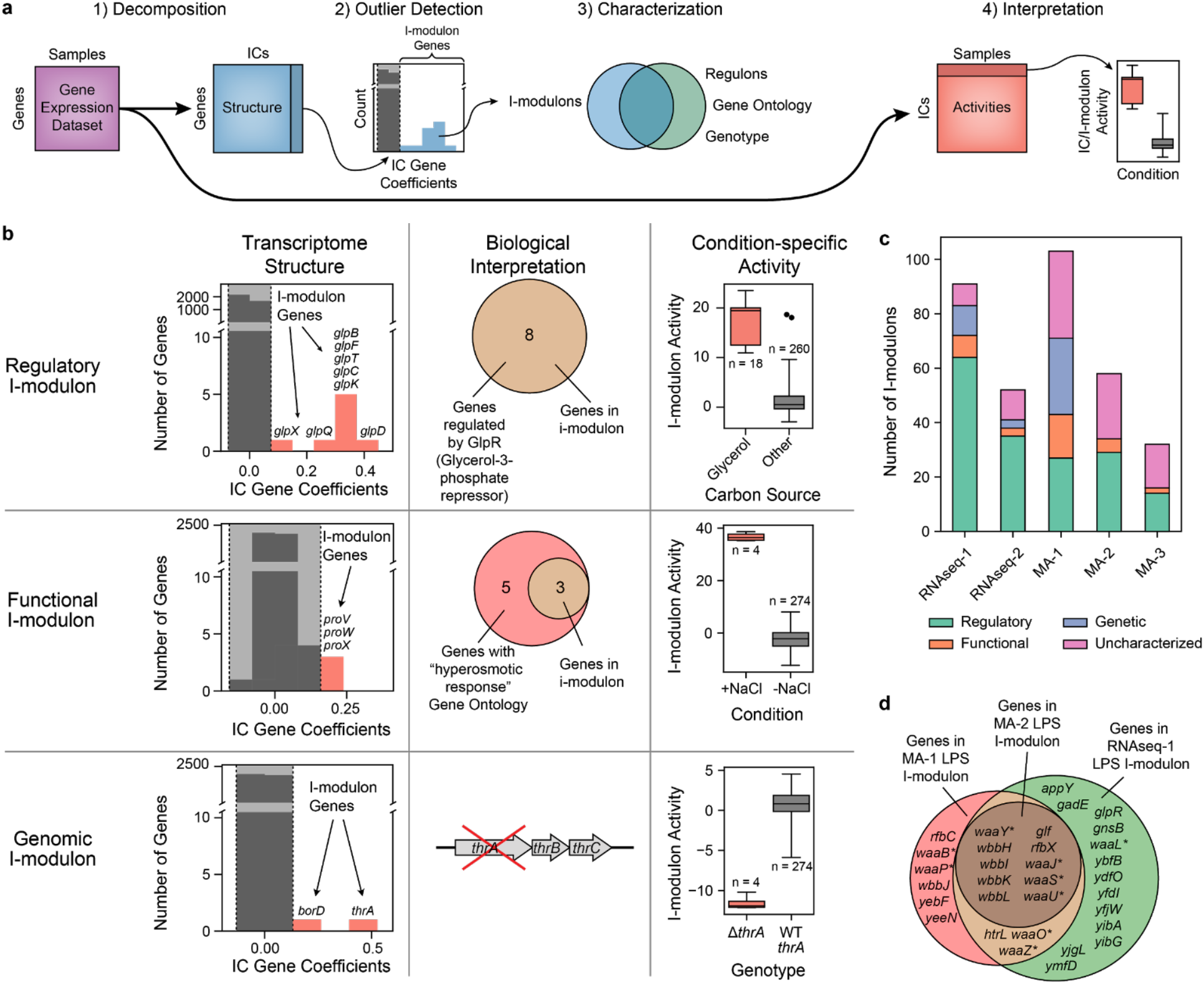
Characterization of i-modulons derived from five independent gene expression datasets. (a) Schematic illustration of the workflow applied to each data set. (b) Descriptions of the three classes of characterized i-modulons. The first column contains histograms illustrating the distribution of gene coefficients in each of three independent components (ICs) from the RNAseq-1 dataset. Genes outside of a threshold (in red) belong to an “i-modulon”. The second column illustrates the biological interpretation of the i-modulon types. I-modulons are characterized by comparing their genes with known regulons, ontological annotations, and genotypes. The third column displays the ICA-computed activity levels for the selected ICs across all 278 conditions in the RNAseq-1 dataset. (c) Bar chart describing the four types of i-modulons computed from each of the five gene expression datasets. (d) Comparison of three Functional i-modulons, each derived from decomposition of a different dataset. Asterisks indicate genes annotated with the GO term “lipopolysaccharide core region biosynthetic process”.

We categorized the resulting i-modulons from each dataset into four classes, based on their gene content: (1) Regulatory, (2) Functional, (3) Genomic, and (4) Uncharacterized (Figure 1b,c, Supplementary Data 4). Prior work showed that i-modulons are highly consistent with, but not identical to, known regulons. On average, these i-modulons captured 80% of the known targets of their linked transcriptional regulators and have accurately predicted new regulon members^30^. Although i-modulons often contain genes known to be regulated by a single transcriptional regulator (e.g. transcription factor, sigma factor, or riboswitch), it has been observed that i-modulons may represent combinations of multiple regulons^19,30^.

In this study, a *Regulatory* i-modulon was defined as an i-modulon that was statistically enriched with a single regulon (Fisher’s exact test, FDR < 10^−5^). Regulatory i-modulons were named after the single transcriptional regulator whose targets provided the best overlap (See Methods).

*Functional* i-modulons contained genes with highly similar functions but lacked common regulator(s). For example, three i-modulons were identified in MA-1, MA-2, and RNAseq-1, respectively, that share 10 genes and were enriched in the gene ontology (GO) term “lipopolysaccharide core region biosynthetic process” (Figure 1d). These i-modulons were named based on the GO term with the lowest enrichment p-value (FDR < .01). This category also included i-modulons composed of prophages with no known regulators. Functional i-modulons represent a compelling opportunity for discovery of new transcriptional regulators^41^.

*Genomic* i-modulons reflect alterations in the genome, such as those resulting from engineered overexpression or knock-out of one or more genes. The activities of these i-modulons represent the presence of these genomic alterations, such as large-scale duplications or deletions. These i-modulons can be useful to validate strain-specific mutations or deletions^30^ and successful plasmid transformations^42^. Genomic i-modulons were categorized by reconciling i-modulon genes and activities with strain-specific genotypes.

The remaining *Uncharacterized* i-modulons could not be interpreted, either due to a high number of uncharacterized genes or the presence of seemingly functionally unrelated genes. Uncharacterized i-modulons may represent undiscovered regulons, noise, or other unwanted sources of variation in the datasets.

The RNA-seq datasets produced the highest fraction of characterized i-modulons; only 9% of i-modulons in RNAseq-1, and 21% of i-modulons in RNAseq-2 were Uncharacterized. In contrast, we were unable to characterize 30-50% of the i-modulons derived from the microarray compendia. The largest fraction of uncharacterized i-modulons was observed in MA-3, which also contained the fewest samples.

### The i-modulon structure is conserved across transcriptomic datasets

Each of the five transcriptomic datasets were created using different technologies or generated by different research groups, spanning fifteen years (2004-2019). In addition, each dataset contained vastly different experimental conditions and genotypes, such as overexpressed cellular division proteins in the MA-1 dataset^32^, diverse nutritional supplements in the RNAseq-1 dataset, or strains evolved to resist antibiotics in the RNAseq-2 dataset^31^. Although the presence of different conditions in each dataset complicated the identification of consistent i-modulons, many of the i-modulons generated from the five datasets unexpectedly shared similar genes and annotations.

For example, each dataset produced an i-modulon enriched in the sulfur utilization regulator CysB. The IC gene coefficients between four of these i-modulons were highly correlated (Pearson R > 0.5), indicating that the genes in the i-modulons were modulated at similar ratios across all five datasets regardless of expression profiling platform (Figure 2a). However, the CysB i-modulon from the MA-3 dataset exhibited poorer correlations with the other i-modulons. The MA-3 CysB i-modulon contained many genes that were absent from any other CysB i-modulon. Many of these genes were involved in the biosynthesis of other amino acids and were present in different i-modulons in the other datasets. In fact, the MA-3 CysB i-modulon could be approximated (R^2^ = 0.47) by a linear combination of 10 i-modulons from the RNAseq-1 dataset related to nutrient availability (Figure 2b, Supplementary Table 1). This example demonstrates that an i-modulon and its IC gene coefficients are unaffected by technology or dataset composition, given a diverse enough dataset.

**Figure 2:**
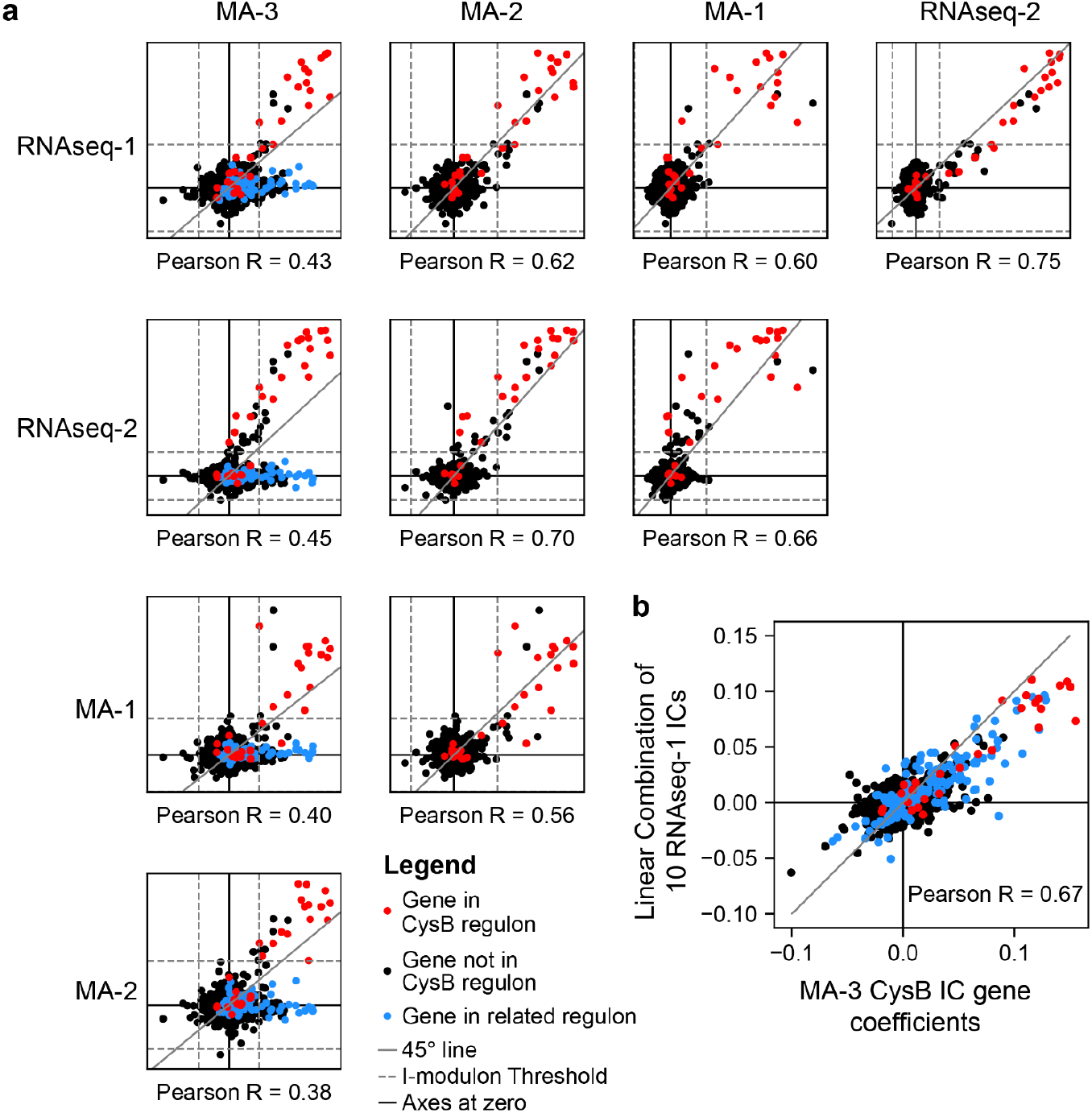
Comparison of IC gene coefficients for i-modulons enriched with genes in the CysB regulon. (a) Scatter plot between IC gene coefficients for CysB-linked i-modulons across all five datasets. Genes in the published CysB regulon are colored in red. For comparisons in the first column involving the MA-3 dataset, genes regulated by any of the following regulators are colored in blue: MetJ, TrpR, GlpR,ArgR, Lrp, CysB, leu-tRNA-mediated transcriptional attenuation, or thiamine riboswitch. All other genes are black. Dashed lines indicate i-modulon thresholds. Gray solid lines indicate the 45-degree line of equal gene coefficients. (b) Scatter plot between IC gene coefficients for the CysB i-modulon in MA-3 compared to a linear combination of 10 ICs from RNAseq-1 (see Supplementary Table 1). Color scheme is identical to panel (a).

To extend our assessment of i-modulons reproducibility, we compared all i-modulons found in each dataset using a Reciprocal Best Hit (RBH) graph^29^ (Figure 3a, Supplementary Table 2). In the RBH graph, each node represents an i-modulon, and nodes are connected when i-modulons from two different datasets find each other as the best scoring i-modulon in the other dataset^43^. I-modulons were scored by the absolute correlation between the gene coefficients in their respective ICs, as shown for the CysB i-modulon cluster. For clarity, edges with low scores were removed from the graph (See methods, Supplementary Figure 1).

**Figure 3:**
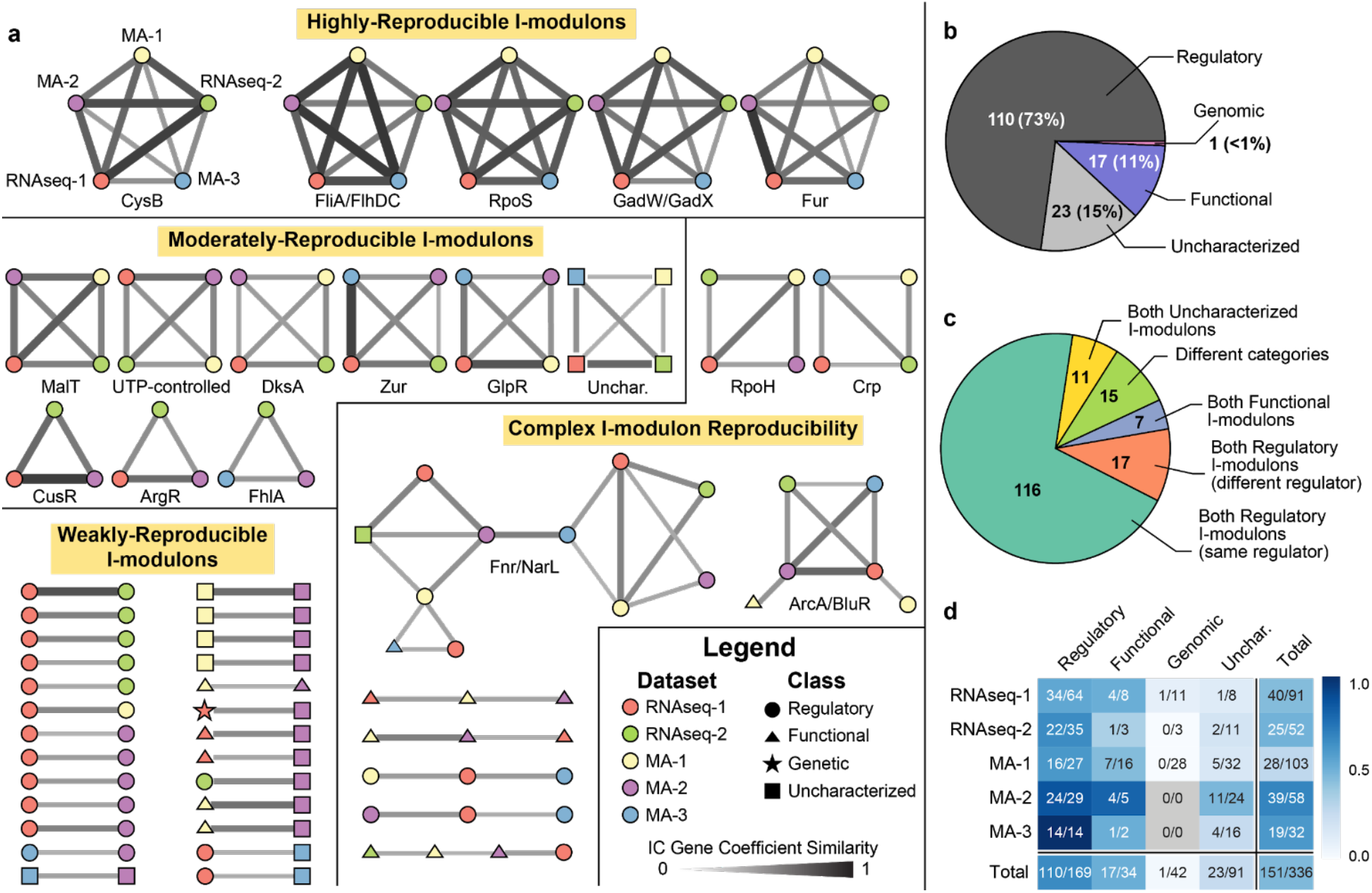
The i-modulon structure is conserved across five datasets. (a) Reciprocal best hit (RBH) graph indicating i-modulons as nodes and RBHs as edges. Node color indicates the source dataset for the i-modulon, as denoted for the CysB i-modulon. Node shape indicates the i-modulon category. Edge thickness and darkness indicate the gene coefficient similarity. Clusters are labelled with the regulator(s) that are linked to the i-modulons in the cluster, if available, and grouped by level of i-modulon reproducibility.Pie chart describing the categories of all i-modulons shown in the RBH graph. (c) Pie chart describing the types of edges in the RBH graph. (d) Heatmap indicating how many i-modulons from each dataset and category were in the RBH graph.

Of the 336 i-modulons identified across all five datasets, nearly half (45%) were linked to an i-modulon in another dataset. Of these 151 reproducible i-modulons, 110 (73%) were classified as Regulatory, and the remaining were either Functional or Uncharacterized (Figure 3b). Only one Genomic i-modulon was matched to an i-modulon from another dataset (Supplementary Figure 2).

Most edges in Figure 3a connected two Regulatory i-modulons enriched in the same regulator (Figure 3c). Across all five datasets, Regulatory i-modulons were the most reproducible category (Figure 3d). The lowest fraction of reproducible Regulatory i-modulons was observed in the RNAseq-1 dataset, likely due to niche transcription factors that are active under specific environmental conditions.

Many i-modulons in the RBH graph separated into well-defined clusters, each linking at most one i-modulon from each dataset. I-modulons in a cluster were often linked to the same regulator, as labelled below each cluster. Five clusters were highly-reproducible, each containing one i-modulon from each of the five datasets. Three of these clusters were enriched with genes from a single regulon, whereas the other two clusters contained i-modulons enriched with genes regulated by closely-related transcriptional regulators. An additional nine i-modulons were found in three or four datasets, forming moderately-reproducible clusters.

However, some clusters exhibited complex connectivity, the largest of which contained i-modulons linked to either of the nitrate response regulators Fnr and NarL. Cellular respiration is regulated by a combination of highly-interconnected transcription factors^33^, indicating that these i-modulons may represent combined effects of regulators (Supplementary Figure 3).

The analysis presented in this section shows that ICA recovers consistent, technology-independent signals across multiple gene expression datasets. This property is unique to ICA as compared to other dimensionality reduction algorithms (Supplementary Figure 4).

### Data integration reveals increased resolution and novel insights

I-modulons are a linear approximation of the structure underlying transcriptomics datasets. Since elements of this structure were highly consistent across the five transcriptomics datasets, we combined the datasets together to explore whether ICA would identify a similar structure in the combined compendium, which contained 802 expression profiles (Figure 4a).

**Figure 4:**
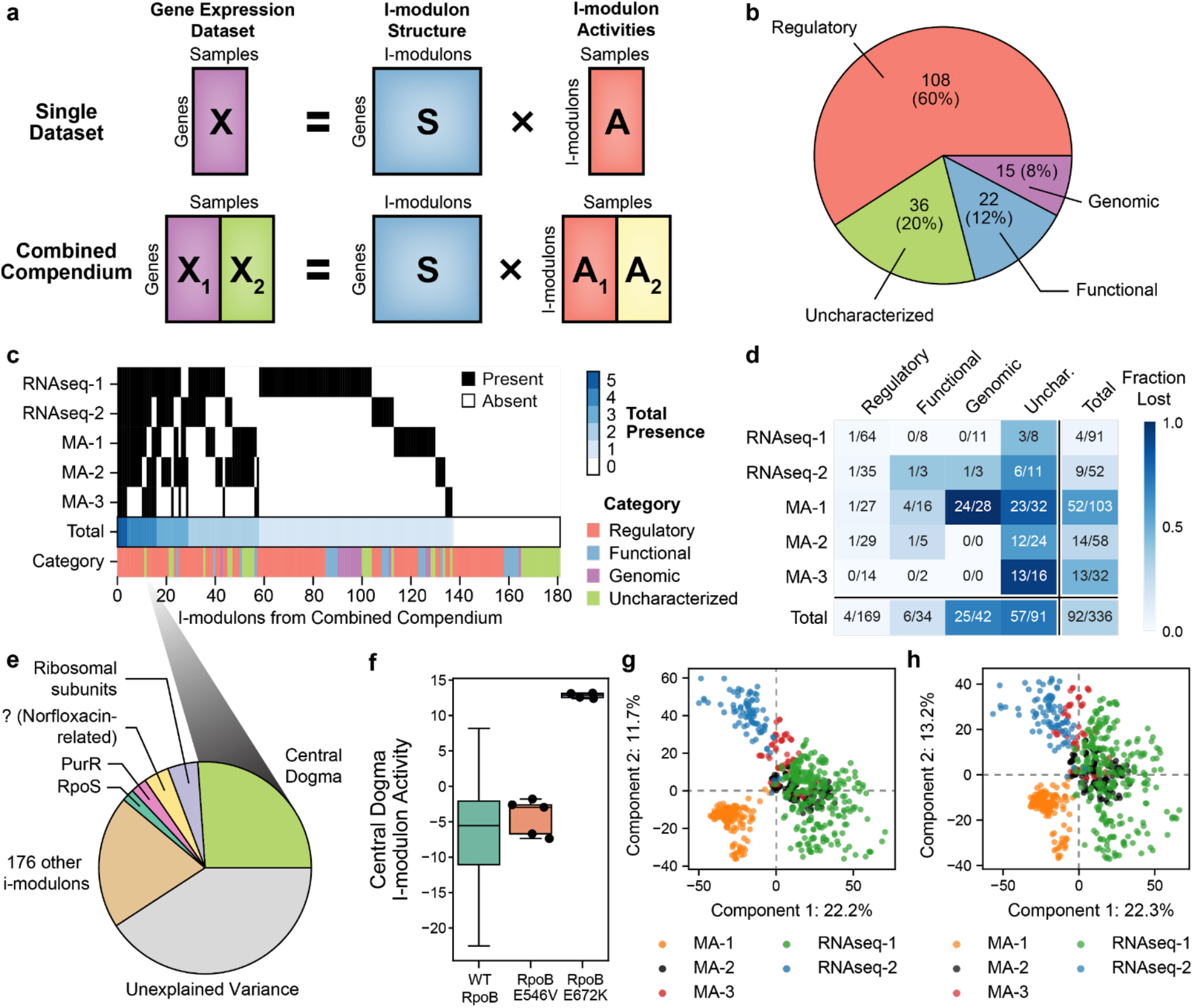
Decomposition of combined compendium results in increased resolution. (a) Schematic illustration of data integration. (b) Pie chart showing the categories of the 181 i-modulons from the combined dataset. (c) Heatmap illustrating which i-modulons from the combined compendium were matched to i-modulons in individual datasets in the RBH graph. The total number of matches for each i-modulon is shown in blue. Category of each i-modulon is shown below. (d) Heatmap illustrating how many i-modulons from each category and dataset were matched to an i-modulon in the full compendium in the RBH graph. (e) Pie chart illustrating the fraction of expression difference resulting from ppGpp-RNAP binding explained by each i-modulon. The norfloxacin-related i-modulon is described in Supplementary Figure 5e. (f) Boxplot of the Central Dogma i-modulon activities for expression profiles in the RNAseq-1 dataset. (g) Principal component loadings of the combined compendium expression levels without batch correction. (h) Principal component loadings of the i-modulon activities from the combined compendium.

We extracted 181 i-modulons from the combined compendium and characterized them as described previously (Figure 4b). The majority of i-modulons from this combined compendium (60%) were enriched with targets of a known transcriptional regulator. To understand the effects of dataset integration, we compared the new i-modulons to the i-modulons from the independent datasets.

First, we asked whether the i-modulons identified in the original datasets were retained upon data integration. From the RBH graph, 75% of the 181 i-modulons in the combined compendium could be directly linked back to at least one i-modulon derived from an individual dataset, many of which were linked to i-modulons from two or more datasets (Figure 4c, Supplementary Table 3). The two datasets with the largest number of unique conditions, (RNAseq-1 and MA-1) contained the most uniquely mapped i-modulons, as these datasets were more likely to activate niche transcriptional regulators.

We next investigated the properties of i-modulons that were not retained upon data integration. Nearly all of the 92 missing i-modulons were categorized as either Genomic or Uncharacterized (Figure 4d). Missing i-modulons were weaker signals, each of which accounted for a significantly lower fraction of expression variance than retained i-modulons (Mann-Whitney-U Test p-value < 10^−5^, Supplementary Figure 5a).

Additionally, 44 new i-modulons in the combined compendium could not be traced back to the i-modulons from the individual datasets. Of these new 44 i-modulons, 21 represented the effects of transcriptional regulators that could not be discriminated in any of the individual datasets alone and 12 were dominated by a single gene (Supplementary Figure 5b,c). A previous study showed that over-decomposition of a transcriptomic dataset results in smaller groups of genes found in each i-modulon but does not affect the biological relevance of other i-modulons^44^.

One Uncharacterized i-modulon in the combined compendium was linked back to four of the five individual datasets. These four i-modulons created the only uncharacterized cluster in Figure 3a. Although this i-modulon was not enriched in any GO term, it contained many genes encoding functions related to the central dogma of molecular biology, such as rRNA and tRNA modification, RNases, and helicases (Supplementary Figure 5d). Therefore, we named it the “Central Dogma” i-modulon. Over half of the genes in this i-modulon are not known to be regulated by any transcription factor, and the remaining genes are not enriched in any common regulator.

A recent study found that binding of ppGpp, the stringent response alarmone^45^, to RNA polymerase (RNAP) directly down-regulated 428 genes^46^, including 48 of the 54 genes in Central Dogma i-modulon. The Central Dogma i-modulon explained 26% of the expression variation between strains affected by ppGpp-RNAP binding and the wild-type strain (Figure 4e). Additionally, an RNAP mutant strain exhibits the highest Central Dogma activity in the RNAseq-1 dataset (Figure 4f), providing evidence that the point mutation affects ppGpp binding to RNAP^47^.

Next, we investigated whether dataset integration could change the quality of the Regulatory i-modulons. Since a Regulatory i-modulon consists of genes in a known regulon, the quality of a Regulatory i-modulon can be assessed using the F1-score, which is the harmonic mean of precision and recall of the i-modulon compared to the regulon (Supplementary Figure 5f). We inspected all Regulatory i-modulons that were found in both an individual dataset and the combined compendium and found that the average F1-score increased from 0.54 to 0.59 (Wilcoxon Rank Sum p-value = 10^−5^) (Supplementary Figure 5g). This result showed that on average, Regulatory i-modulons from the combined compendium were more similar to the previously known regulons than the Regulatory i-modulons from individual datasets.

Finally, we checked whether the i-modulon activities were affected by dataset integration. When we apply ICA to an expression profiling dataset, we obtain the **S** matrix, encoding the i-modulon structure, and an **A** matrix, which contains activity levels for each i-modulon across every expression profile (Figure 1a and Figure 4a). We have focused our analysis thus far on the invariant properties of the **S** matrix, and briefly discuss the effects of data integration on the **A** matrix.

I-modulon activities reflect the overall change in expression of the i-modulon genes and serve as a proxy for transcription factor activities^26^. Therefore, it is important to ensure that the relative i-modulon activities are unchanged upon dataset integration. The absolute Pearson R correlation coefficient between the i-modulon activities of linked components showed that most i-modulon activities are unaffected by dataset integration (Supplementary Figure 5h).

Previously, principal component analysis (PCA) of integrated expression datasets revealed that the source of each dataset was the dominant discriminator ^48,49^. This result is recapitulated in our combined compendium (Figure 4g). However, since the i-modulon structure of each dataset (encoded in **S**) is consistent across datasets, we see that technical heterogeneity is stored in the activity matrix (**A**) (Figure 4h). Therefore i-modulon activities cannot be compared across datasets but can still reliably be compared within the same dataset.

### ICA elucidates similar structures in the *E. coli* proteome

Since the i-modulon structure was highly consistent across transcriptomics datasets, we asked whether shadows of the transcriptional regulatory program could be observed in the proteome. Thus, we applied the extended ICA algorithm on two high-quality proteomics datasets from two independent research groups^50,51^ (Supplementary Datasets 1-3). The Proteomics-1 dataset had higher gene coverage whereas the Proteomics-2 dataset included more conditions (Figure 5a,b).

**Figure 5:**
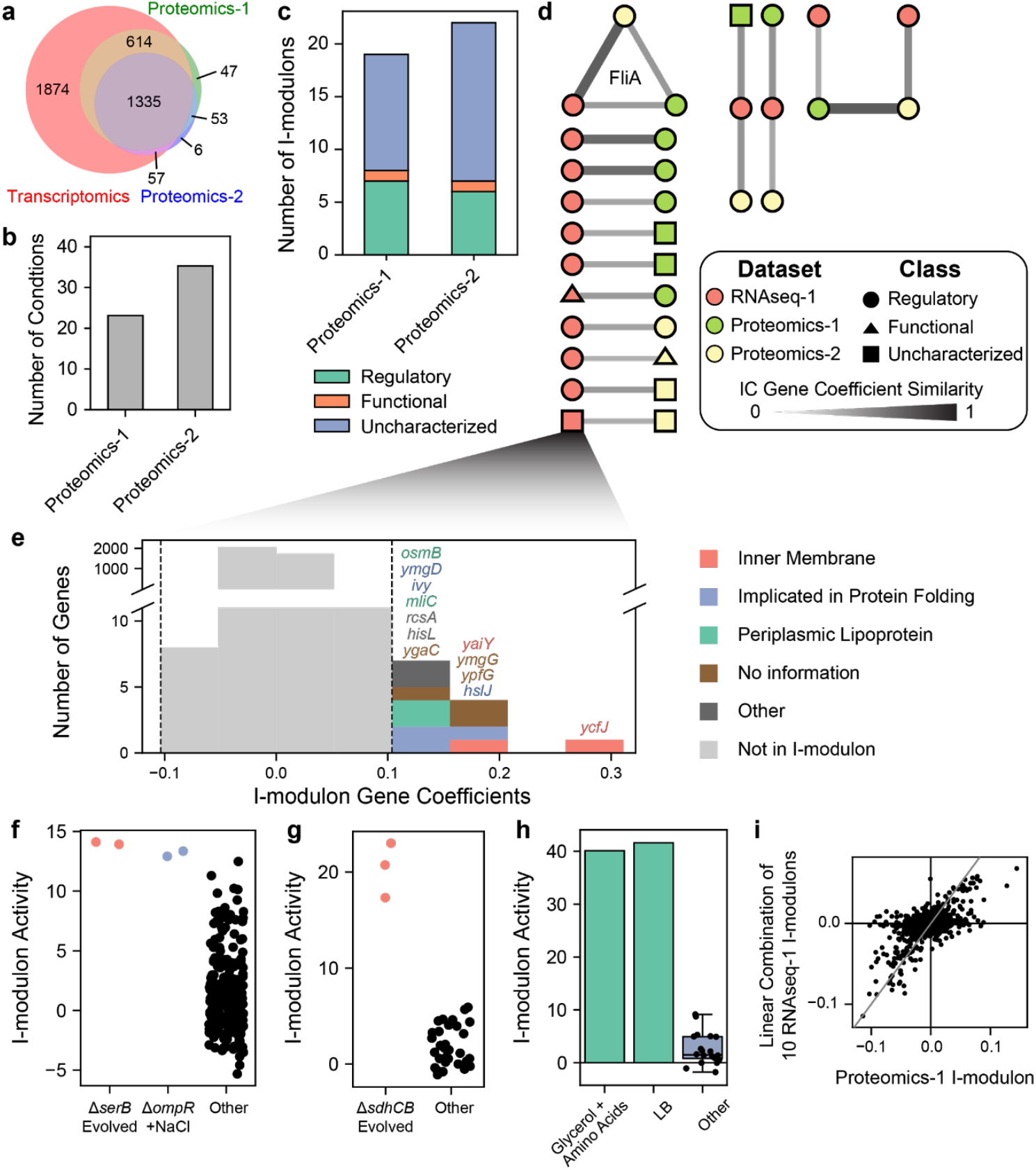
ICA elucidates similar structures in the *E. coli* transcriptome and proteome. (a) Venn diagram showing the coverage of genes in transcriptomics and proteomics datasets. (b) Bar chart of the number of samples in each proteomics dataset. (c) Bar chart showing the types of i-modulons extracted from two independent proteomic datasets. (d) RBH graph between i-modulons from two proteomics datasets and the RNAseq-1 i-modulons. (e) Genes in the RNAseq-1 i-modulon that was also detected in the Proteomics-1 dataset (f) Activity of the Uncharacterized i-modulon illustrated in Panel C. (g) Activity of the Proteomics-2 i-modulon linked to the RNAseq-1 i-modulon illustrated in Panel C (h) Activity levels of an Uncharacterized i-modulon in the Proteomics-1 dataset. (i) Scatter plot between IC protein coefficients for the i-modulon described in panel H compared to a linear combination of 10 ICs from RNAseq-1 (see Supplementary Table 5). Black lines indicate zero IC gene coefficients, and gray line indicates 45-degree line.

We characterized the i-modulons from the proteomics datasets as described previously (Figure 5c, Supplementary Dataset 4), and used the RBH method to compare the proteomic i-modulons to i-modulons from the RNAseq-1 dataset (Figure 5d, Supplementary Table 4). We note that most i-modulons from the proteomics datasets were Uncharacterized, likely due to the intrinsically lower coverage and small dataset sizes.

One pair of Uncharacterized i-modulons were independently detected in the RNAseq-1 dataset and the Proteomics-1 dataset (Supplementary Figure 6). The RNAseq-1 i-modulon contained many membrane-associated genes (Figure 5e) and was active under osmotic stress and in laboratory evolved strains (Figure 5f,g). In addition, many genes in this i-modulon had been identified in two prior studies as genes upregulated by the envelope stress Rcs-phosphorelay^52^ or upregulated in response to beta-lactam antibiotics that inhibit cell wall biosynthesis^53^. Therefore, we propose that all the genes in this i-modulon are co-regulated by a transcription factor related to the Rcs system that may currently be uncharacterized. This represents a clear potential for discovery.

It is important to note that the RNAseq-1 i-modulons were more similar to the other transcriptomics i-modulons than proteomics i-modulons (Mann-Whitney U Test, p-value = 0.005). Although some of this deviation may be due to differences between relative abundances between transcripts and proteins, the small sample sizes of the proteomics datasets resulted in i-modulon coalescence as previously described with the MA-3 CysB i-modulon. In particular, one Uncharacterized i-modulon from the Proteomics-1 dataset was only active under two media conditions: LB rich media, and Glycerol M9 media supplemented with amino acids (Figure 5h). This proteomic i-modulon can be partially approximated (R^2^ = 0.33) by a linear combination of 10 RNAseq-1 i-modulons, (Figure 5i) related to nutrient availability (Supplementary Table 5). Six of these i-modulons coincide with the 10 RNAseq-1 i-modulons comprising the MA-3 CysB i-modulon.

Due to the lower coverage of genes and low number of conditions in the proteomics datasets, it is not currently possible to conclusively state that the i-modulon structure is conserved between proteomics and transcriptomics datasets, but this section provides evidence that such structure exists.

## Discussion

Matrix decomposition is a powerful approach to extract knowledge from large transcriptomics datasets. In particular, we have shown that ICA identifies highly similar structures between dissimilar datasets for the model bacteria *E. coli*. In addition, a combined compendium produced many identical i-modulons as the individual datasets and could distinguish further signals that could not be identified in the separate datasets. The i-modulons derived from the compendium can be applied to interpret new datasets, accelerating discovery and providing a standard framework that could be used to investigate any transcriptional regulator.

Throughout this study, we observed various properties of the i-modulon decompositions:(1) i-modulons co-occurring in multiple datasets tend to represent the effects of transcriptional regulators, (2) i-modulon detection from a data set is dependent on the experimental conditions used to generate it, (3) transcriptomic and proteomic i-modulons are surprisingly similar, (4) ICA can be applied to cross-platform transcriptomic compendia without the need for normalization, and (5) integration of data sets reveals i-modulons not found in individual data sets.

In all five transcriptomic *E. coli* datasets, most i-modulons could be characterized as representing the effects of a transcriptional regulator or a gene knock-out (i.e. Regulatory or Genomic). However, when extending to less-characterized organisms, it could be difficult to determine whether the remaining i-modulons are technical artifacts or contain true biological insight, as demonstrated by the Central Dogma i-modulon. This example demonstrates that if an i-modulon is identified in multiple datasets, or if an i-modulon persists upon addition of new datasets, then it could represent a true transcriptional signal.

Another important note is that the i-modulons extracted from each dataset are sensitive to the experimental conditions that are represented in the dataset; ICA cannot extract an i-modulon for a transcription factor whose activity never changes. Additionally, the CysB i-modulons demonstrated that i-modulons may represent the effects of multiple regulators, when the activities of the regulators are highly correlated across the measured conditions. However, adding new data or increasing the dimensionality of the decomposition can decouple the regulators, splitting the i-modulon into its biological parts^30,44^.

In contrast to the condition-invariant i-modulon structure, i-modulon activities represent the condition-dependent dynamics of expression profiles. In this study, we do not apply any normalization techniques to the data and clearly observe batch effects in the activity matrix of the combined compendium. Further work comparing identical experimental conditions from separate protocols is required to enable i-modulon activity comparisons across datasets.

We have shown that the *E. coli* transcriptome contains a conserved, underlying structure that is found across multiple independent datasets and extends into the proteome. Previously, we also showed that this structure also exists across strains within a species^30^. This powerful observation enables unprecedented re-analyses of thousands of previously published datasets and demonstrates how data science can unlock hidden potential in complex biological datasets.

## Methods

### RNA-seq and Microarray Data Processing

The full, log-transformed transcripts-per-million (log-TPM) for the RNAseq-1 dataset^30^ was obtained from https://github.com/SBRG/precise-db. Raw data comprising the RNAseq-2 compendium were obtained from NCBI SRA under the bioproject accession number PRJNA379428^31^. Raw data was processed using a similar process as described in Sastry et al. 30. Raw sequencing reads were mapped to the reference genome (NC_000913.3) using bowtie (v1.1.2)^54^ with the following options “-X 1000 -n 2 -3 3”. Transcript abundance was quantified using summarizeOverlaps from the R GenomicAlignments package (v1.18.0), with the following options “mode = “IntersectionStrict”, singleEnd = FALSE, ignore.strand = FALSE, preprocess.reads = invertStrand”^55^. Transcripts per million (TPM) were calculated by DESeq2 (v1.22.1)^56^. The final expression dataset was log-transformed log_2_(TPM + 1) before analysis, referred to as log-TPM.

CEL files were obtained for the three microarray datasets from NCBI GEO^1^ (see Supplementary Data 1 for accession numbers). In order to build large enough datasets for analysis, datasets MA-2 and MA-3 included all public data available from our research group that used the same expression platform. Each microarray dataset was normalized using robust multichip average (RMA) with default arguments from the R package *affy*^57^.

As the probes in microarrays may vary across platforms, only genes that were measured in all 5 datasets (3880 genes) were included in the analysis. Low quality expression profiles that clustered separately from the rest of the dataset were removed from the MA-1 and MA-2 datasets (See Supplementary Data 1). Each dataset was individually centered by subtracting the average expression profile across the replicates of a dataset-specific reference condition (see Supplementary Data 1). To create the combined compendium, the centered datasets were concatenated, without any additional normalization.

Only genes measured in all five datasets were retained, resulting in 3880 genes per dataset. Gene names, b-numbers, operons, and descriptions were obtained from Ecocyc^58^. Clusters of orthologous groups (COG) annotations were obtained from eggNOG 4.5.1^59^. Additional annotations were obtained from Gene Ontology^60^. The transcriptional regulatory network (TRN) for transcription factors, small RNAs, and sigma factors was obtained from RegulonDB v10.0^61^, and supplemented with newly identified regulons from recent publications^30,41^. Riboswitches, tRNA-mediated attenuation, and dksA binding sites were obtained from Ecocyc^58^, and UTP/CTP-dependent attenuation and reiterative transcription were obtained from Turnbough and Switzer^62^. All annotations for the 3880 genes are reported in Supplementary Dataset 5.

### Proteomics data processing

Processed data for the Proteomics-1 dataset was acquired from Schmidt et al.^50^, using the values reported for protein copies per cell. Only genes with measurements across all samples were retained, resulting in 2058 genes. Processed data for the Proteomics-2 dataset was acquired from Heckmann et al.^51^ and augmented with additional samples for growth on different carbon sources that were acquired as described in Heckmann et al.^51^ (See Supplementary Dataset 1 for details). Protein abundance estimation is described in Heckmann et al.^51^; in short, the top3 metric^63,64^ was calibrated using the UPS2 standard to obtain protein amount loaded based on the average of three technical replicates per sample. For each strain, two biological replicates were profiled, and abundances were averaged before applying ICA.

### Independent Component Analysis

ICA decomposes a matrix (**X**) into two matrices: **S** contains the independent signals, or structure, of the dataset, and **A** contains the condition-dependent activities of the signals:

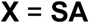

ICA was applied to each individual dataset and the combined compendium, as described in Sastry et al.^30^. Briefly, we executed FastICA 100 times with random seeds and a convergence tolerance of 10^−6^ for RNA-seq data, and a convergence tolerance of 10^−7^ for proteomics data. We constrained the number of independent components (ICs) in each iteration to the number of components that reconstruct 99% of the variance as calculated by principal component analysis. The resulting ICs were clustered using DBSCAN to identify robust ICs, with parameters with epsilon of 0.1, and minimum cluster seed size of 50. This process was repeated 10 times, and only ICs that consistently occurred in all runs were kept.

As described in Sastry et al.^30^, i-modulons were extracted from ICs by iteratively removing genes with the largest absolute value and computing the D’Agostino K^2^ test statistic of the resulting distribution. Once the test statistic fell below a cutoff, we designated the removed genes as the “i-modulon”.

To identify this cutoff for each individual dataset, we performed a sensitivity analysis on the concordance between significant genes in each IC and all known regulons. First, we isolated the 20 genes from each IC (10 genes for proteomics i-modulons) with the highest absolute gene coefficients. We then compared each gene set against all known regulons using the one-sided Fisher’s Exact Test (FDR < 10^−5^). For each component with at least one significant enrichment, we selected the regulator with the lowest p-value.

Next, we varied the D’Agostino K^2^ test statistic from 200 through 1000 in increments of 50. Using the protocol defined above, i-modulons were extracted from ICs at each test statistic value, and the F1-score (harmonic average between precision and recall) was computed between the significant genes and its linked regulator. The test statistic with the maximum F1-score was used as the test statistic cutoff for the respective dataset.

### Characterizing I-modulons

We compared the set of significant genes in each i-modulon to each regulon (defined as the set of genes regulated by any given regulator) using the one-sided Fisher’s Exact Test (FDR < 10^−5^). We then compared the significant genes in each i-modulon to the genes in each Gene Ontology (GO) term using the one-sided Fisher’s Exact Test (FDR < .01). We added prophage information from Ecocyc to our GO database to capture i-modulons representing prophages. In general, the final annotation was selected by the enrichment with the lowest q-value. Some i-modulon annotations were manually curated, as denoted in Supplementary Dataset 5. Genomic i-modulons were also manually curated by (1) comparing i-modulon genes to known genetic alterations (e.g. knock-outs or overexpression), and (2) validating that the i-modulon activities were affected in the appropriate direction for the corresponding strain.

To account for the lower coverage of the proteomics datasets, we trimmed the TRN and GO databases to only include relevant genes. We compared proteomics i-modulons to regulons with FDR < 10^−3^ and compared proteomics i-modulons to GO terms with FDR < .01.

### Comparing I-modulon Structures

To compare the complete structure of the transcriptomic datasets, we constructed the reciprocal best hit (RBH) graphs using the full IC gene coefficients, rather than just the i-modulon genes. We generated the RBH graph as described in Cantini et al. 2019^29^, using the following distance metric to compare ICs:

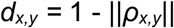

where *ρ_x,y_* is the Pearson correlation between components *x* and *y*.

We pruned all RBHs to remove links between ICs with similarities below 0.3. The full graph is shown in Supplementary Figure 1. The RBH graph was plotted using GraphViz^65^.

### Linear Regression of I-modulons

Regression of the MA-3 CysB i-modulon and the Proteomics-1 i-modulon were performed using the LinearRegression function from Scikit-learn^66^. The ten ICs from the RNAseq-1 dataset with the highest absolute IC gene correlations with each of the CysB i-modulon and Proteomics-1 i-modulon were selected as regression variables for their respective regressions (See Supplementary Tables 1 and 5).

### Inference of I-modulon activities

Raw reads for the ppGpp-RNAP dataset were downloaded from NCBI SRA (PRJNA504613) and processed as described above into log-TPM expression values. Two experimental conditions were selected for comparison, both using the wild-type strain with active *relA*, at 0 and 5 minutes after IPTG induction. Log-TPM expression values were averaged across triplicates.

To infer i-modulon activities and calculate the amount of variance that i-modulons explain between the two datasets, we first centered the two averaged expression profiles, then computed the gene-wise difference. The change in i-modulon activity was calculated by multiplying the expression difference (Δ**X**) by the pseudo-inverse of the S matrix from the full compendium:

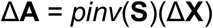

Where *pinv* is the pseudoinverse function.

### Explained Variance

Explained variance between two conditions was calculated as follows:

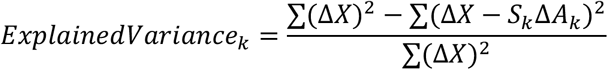

Where *k* is the i-modulon of interest.

## Code availability

Code central to the conclusions is described in the methods and available at https://github.com/SBRG/precise-db. Additional code is available upon request.

## Acknowledgements

The authors would like to thank Dr. Dan Zielinski, Prof. Eivind Almaas and Yara Seif, and CJ Norsigian for informative discussions. This research used resources of the National Energy Research Scientific Computing Center, a DOE Office of Science User Facility supported by the Office of Science of the U.S. Department of Energy under Contract No. DE-AC02-05CH11231. This work was funded by the Novo Nordisk Foundation Center for Biosustainability and the Technical University of Denmark (grant number NNF10CC1016517).

## Author Contributions

A.V.S. and B.O.P. designed the study. A.V.S., A.H., and D.H. performed the analysis. E.K. and S.P. contributed to the interpretation of the data. A.V.S., D.H., E.K., S.P., and B.O.P. wrote and edited the manuscript.

