## Supplemental Information for "Matrix factorization recovers consistent regulatory signals from disparate datasets"

### Supplemental Figures:

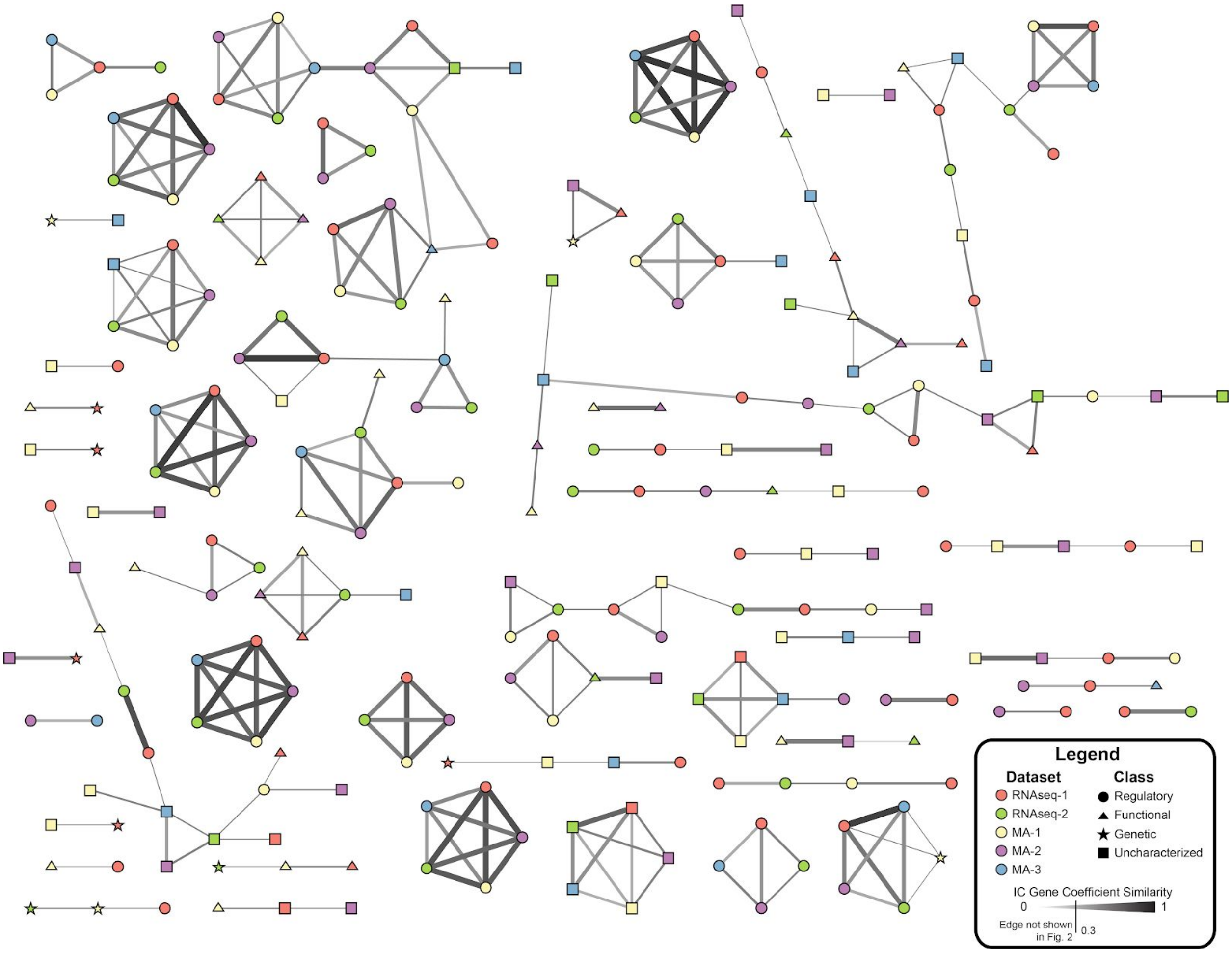


**Supplementary Figure 1: Reciprocal best hit (RBH) graph containing all edges from the five datasets.** Edges with an IC Gene Coefficient similarity score below 0.3 were pruned from Figure 2.

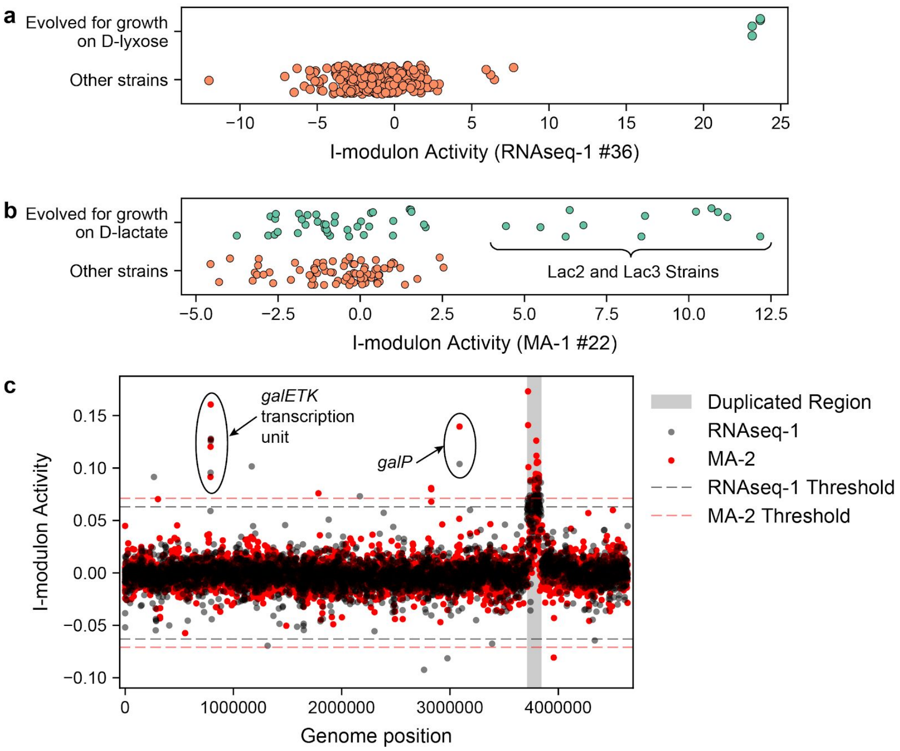


**Supplementary Figure 2: Characteristics of the only Genomic i-modulon found in more than one dataset.** (a) Activities of the i-modulon in the RNAseq-1 dataset separate *E. coli* strains evolved for growth on the non-native carbon source D-lyxose from the other strains in the compendium(Guzmán et al., 2019). (b) Activities of the uncharacterized i-modulon in MA-1 that was linked to the i-modulon described in panel (a). Seven strains were evolved in parallel for growth on D-lactate(Fong et al., 2005), but only two endpoint strains (named Lac2 and Lac3) exhibited high i-modulon activities. These strains were not resequenced, so the adaptive mutations could not be confirmed. (c) Scatter plot showing the IC gene coefficients corresponding to the i-modulons described in panel (a) (in black) and panel (b) (in red). Thresholds determining i-modulon composition are indicated by dashed lines. The genomic duplication from the D-lyxose-evolved strains is highlighted in gray, indicating that all strains with high activities likely acquired an identical duplication along their evolutionary trajectory. Two transcription units outside of the duplicated region were captured in both i-modulons, the *galETK* transcription unit, and the *galP* gene. These genes are responsible for D-galactose catabolism.

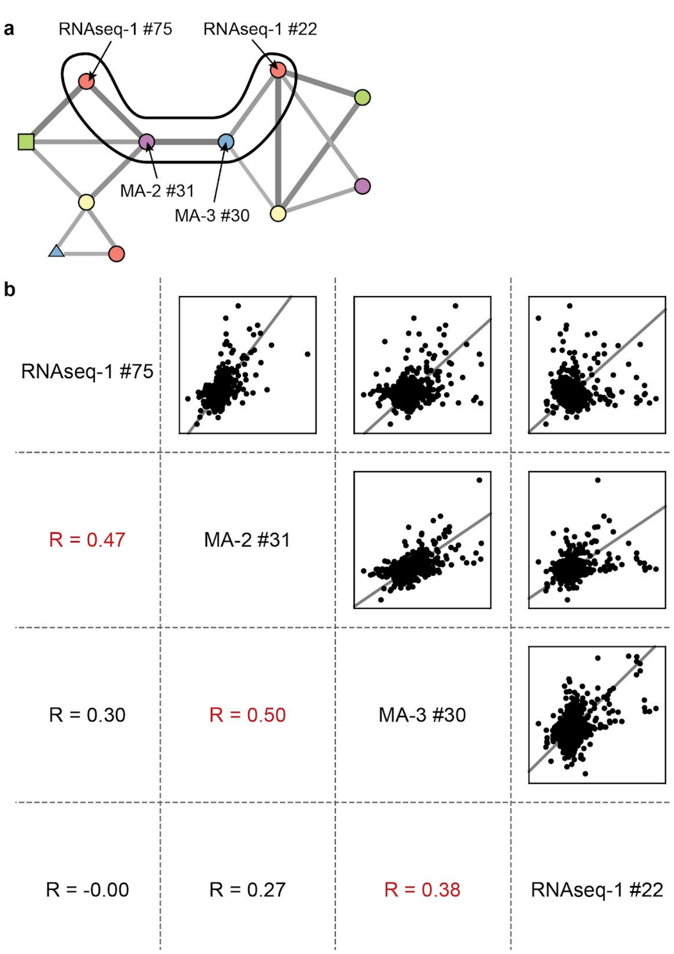


**Supplementary Figure 3: Investigation of a complex i-modulon cluster.** Each dataset contains a different set of conditions, which can activate different groups of respiration-related genes leading to different co-expression patterns between datasets. (a) The complex i-modulon cluster shows that two i-modulons from the RNAseq-1 dataset are indirectly connected. (b) Scatterplots of the IC gene weights for the i-modulons highlighted in (a). The Pearson R correlation of the IC gene weights is shown below. The two RNAseq-1 i-modulons show no correlation, but still contain a few genes in common, indicating that the expression of these shared genes are controlled by two distinct underlying sources.

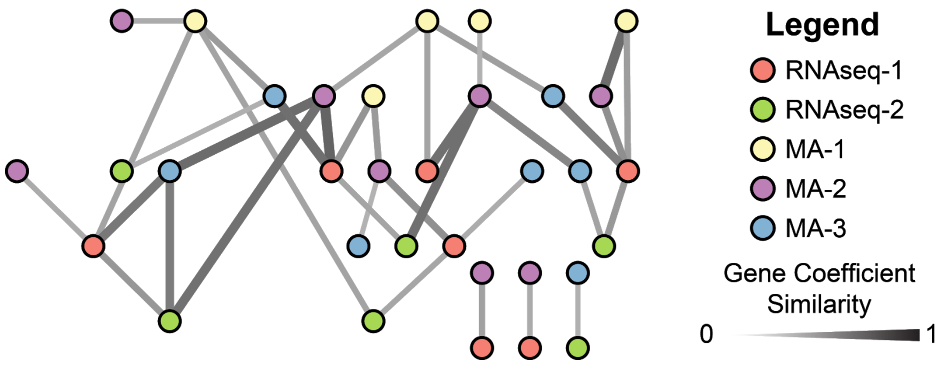


**Supplementary Figure 4: Reciprocal best hit (RBH) graph of Principal Components from all 5 transcriptomic datasets.** Edges with gene coefficient similarity score below 0.3 were pruned from this figure.

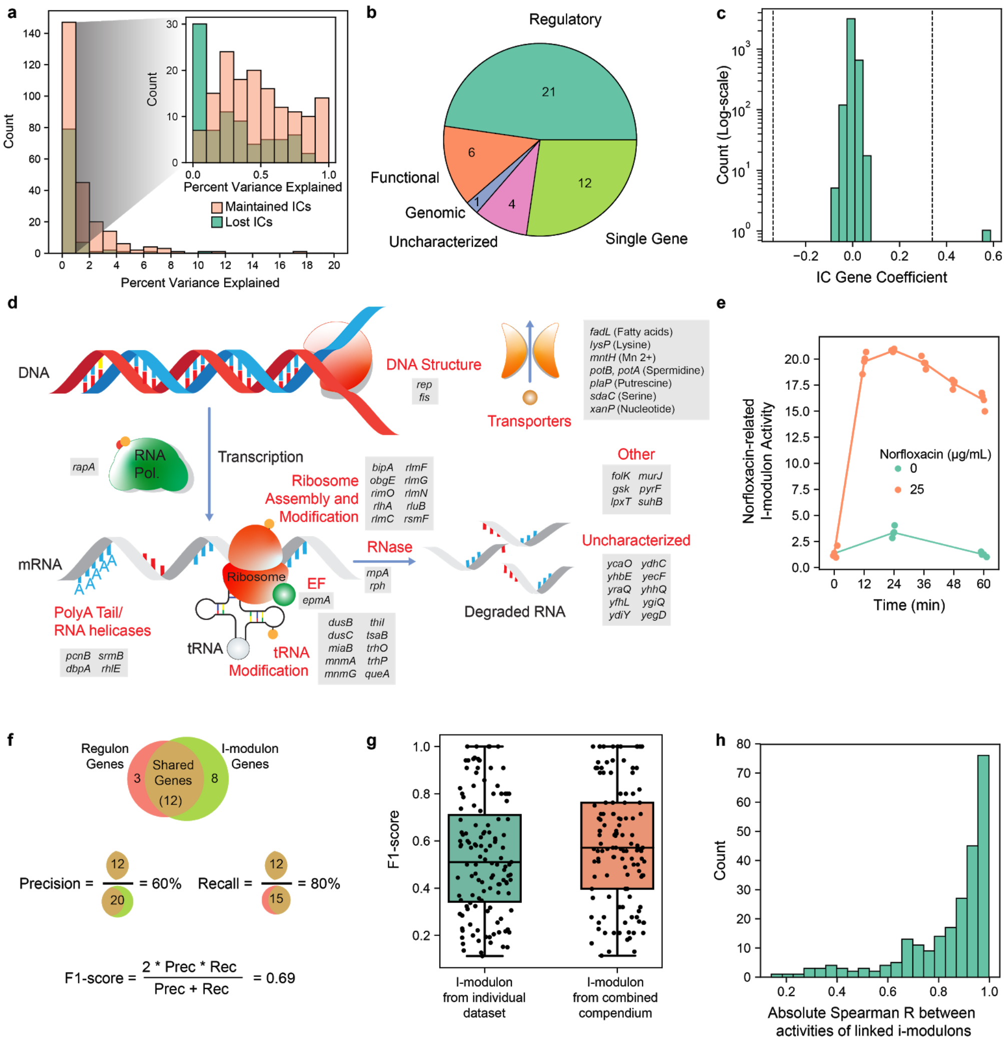

**Supplementary Figure 5: Effects of data integration on i-modulon structure and activity.** (a) Histogram of the percent of total expression explained by each independent component in the individual datasets. Components that are maintained (i.e. identify an RBH hit with the full compendium decomposition) are colored in pink, whereas components that are lost upon data integration are colored in blue. (b) Pie chart illustrating the classes of the new i-modulons extracted from the combined datasets. (c) Histogram of the IC gene coefficients of a Single Gene component. Dashed lines indicate the i-modulon threshold. (d) Schematic illustration of the various processes encoded by the genes in the Central Dogma i-modulon. (e) Time-course treatment of DNA-damage inducing norfloxacin activates an i-modulon that shows reduced activity when RNAP is bound by ppGpp. (f) Schematic illustration of precision, recall, and F1-score. (g) Boxplots of the F1-scores between i-modulons and their associated regulators for Regulatory i-modulons. Only Regulatory i-modulons in the individual dataset that found an RBH in the full compendium are shown in the left boxplot, and the RBH of these i-modulons in the full dataset are shown in the right boxplot. (h) Histogram of absolute Spearman correlations between i-modulon activities in components that are RBHs in the full dataset compared to the individual datasets.

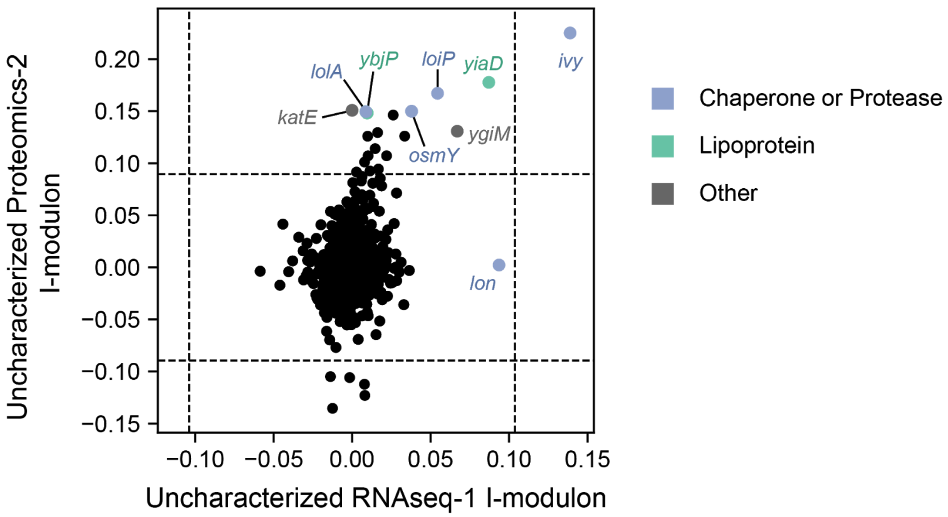


**Supplementary Figure 6: ICA elucidates similar structures from the *E. coli* transcriptome and proteome.** Scatter plot between the two uncharacterized i-modulons highlighted in Figure 4. Although most genes in the RNAseq-1 i-modulon are absent from the proteomics dataset, the genes with the highest IC gene coefficients in either i-modulon are mostly chaperone proteins (mostly periplasmic), proteases, or lipoproteins.

### Supplemental Tables:

**Supplementary Table 1:** Linear regression coefficients for the 10 RNAseq-1 ICs that approximate the MA-3 CysB IC (See Figure 2b)

| **I-modulon** | **Description** | **Linear Regression Coefficient** |
|---|---|---|
| RNAseq-1_46 | CysB | 0.44 |
| RNAseq-1_54 | Thiamine Riboswitch | 0.29 |
| RNAseq-1_17 | ArgR | 0.20 |
| RNAseq-1_87 | leu-tRNA-mediated attenuation | 0.19 |
| RNAseq-1_77 | TyrR | 0.15 |
| RNAseq-1_73 | MetJ | 0.15 |
| RNAseq-1_58 | GlpR | 0.11 |
| RNAseq-1_3 | Cellular response to nutrient levels | -0.11 |
| RNAseq-1_78 | Uncharacterized | -0.14 |
| RNAseq-1_24 | Lrp | -0.20 |

**Supplementary Table 2:** Reciprocal best hits between i-modulons from each transcriptomics dataset. Distances above 0.7 have been pruned.

| I-modulon 1 | I-modulon 2 | Type1 | Type2 | Distance |
|---|---|---|---|---|
| RNAseq-1_0 | RNAseq-2_35 | regulatory | regulatory | 0.47 |
| RNAseq-1_4 | RNAseq-2_37 | regulatory | regulatory | 0.56 |
| RNAseq-1_8 | RNAseq-2_31 | regulatory | regulatory | 0.52 |
| RNAseq-1_10 | RNAseq-2_34 | regulatory | regulatory | 0.5 |
| RNAseq-1_14 | RNAseq-2_27 | regulatory | regulatory | 0.51 |
| RNAseq-1_16 | RNAseq-2_45 | regulatory | regulatory | 0.44 |
| RNAseq-1_17 | RNAseq-2_5 | regulatory | regulatory | 0.56 |
| RNAseq-1_22 | RNAseq-2_7 | regulatory | regulatory | 0.54 |
| RNAseq-1_27 | RNAseq-2_0 | regulatory | regulatory | 0.68 |
| RNAseq-1_33 | RNAseq-2_39 | regulatory | regulatory | 0.33 |
| RNAseq-1_37 | RNAseq-2_47 | regulatory | regulatory | 0.56 |
| RNAseq-1_41 | RNAseq-2_30 | regulatory | regulatory | 0.63 |
| RNAseq-1_44 | RNAseq-2_17 | regulatory | regulatory | 0.66 |
| RNAseq-1_46 | RNAseq-2_25 | regulatory | regulatory | 0.25 |
| RNAseq-1_63 | RNAseq-2_6 | regulatory | regulatory | 0.42 |
| RNAseq-1_64 | RNAseq-2_44 | regulatory | regulatory | 0.29 |
| RNAseq-1_65 | RNAseq-2_24 | regulatory | regulatory | 0.49 |
| RNAseq-1_66 | RNAseq-2_18 | regulatory | regulatory | 0.32 |
| RNAseq-1_75 | RNAseq-2_4 | regulatory | uncharacterized | 0.52 |
| RNAseq-1_77 | RNAseq-2_12 | regulatory | regulatory | 0.65 |
| RNAseq-1_79 | RNAseq-2_8 | uncharacterized | uncharacterized | 0.4 |
| RNAseq-1_83 | RNAseq-2_26 | regulatory | regulatory | 0.53 |
| MA-1_1 | RNAseq-1_82 | regulatory | regulatory | 0.64 |
| MA-1_5 | RNAseq-1_64 | regulatory | regulatory | 0.39 |
| MA-1_6 | RNAseq-1_8 | regulatory | regulatory | 0.59 |
| MA-1_12 | RNAseq-1_46 | regulatory | regulatory | 0.4 |
| MA-1_15 | RNAseq-1_58 | regulatory | regulatory | 0.34 |
| MA-1_16 | RNAseq-1_62 | biological | biological | 0.63 |
| MA-1_34 | RNAseq-1_16 | regulatory | regulatory | 0.44 |
| MA-1_38 | RNAseq-1_37 | regulatory | regulatory | 0.66 |
| MA-1_39 | RNAseq-1_41 | regulatory | regulatory | 0.46 |
| MA-1_49 | RNAseq-1_65 | regulatory | regulatory | 0.26 |
| MA-1_52 | RNAseq-1_10 | regulatory | regulatory | 0.37 |
| MA-1_58 | RNAseq-1_59 | regulatory | regulatory | 0.62 |
| MA-1_68 | RNAseq-1_33 | regulatory | regulatory | 0.33 |
| MA-1_72 | RNAseq-1_22 | regulatory | regulatory | 0.51 |
| MA-1_80 | RNAseq-1_76 | regulatory | regulatory | 0.54 |
| MA-1_82 | RNAseq-1_79 | uncharacterized | uncharacterized | 0.69 |

| | | | | |
|---|---|---|---|---|
| MA-1_89 | RNAseq-1_4 | regulatory | regulatory | 0.58 |
| MA-1_1 | RNAseq-2_4 | regulatory | uncharacterized | 0.65 |
| MA-1_5 | RNAseq-2_44 | regulatory | regulatory | 0.38 |
| MA-1_6 | RNAseq-2_31 | regulatory | regulatory | 0.55 |
| MA-1_10 | RNAseq-2_21 | biological | biological | 0.66 |
| MA-1_12 | RNAseq-2_25 | regulatory | regulatory | 0.34 |
| MA-1_34 | RNAseq-2_45 | regulatory | regulatory | 0.4 |
| MA-1_39 | RNAseq-2_30 | regulatory | regulatory | 0.56 |
| MA-1_49 | RNAseq-2_24 | regulatory | regulatory | 0.48 |
| MA-1_52 | RNAseq-2_34 | regulatory | regulatory | 0.51 |
| MA-1_67 | RNAseq-2_0 | regulatory | regulatory | 0.58 |
| MA-1_68 | RNAseq-2_39 | regulatory | regulatory | 0.36 |
| MA-1_72 | RNAseq-2_7 | regulatory | regulatory | 0.55 |
| MA-1_82 | RNAseq-2_8 | uncharacterized | uncharacterized | 0.67 |
| MA-1_89 | RNAseq-2_37 | regulatory | regulatory | 0.61 |
| MA-1_0 | MA-2_7 | biological | regulatory | 0.58 |
| MA-1_1 | MA-2_31 | regulatory | regulatory | 0.58 |
| MA-1_5 | MA-2_41 | regulatory | regulatory | 0.29 |
| MA-1_6 | MA-2_20 | regulatory | regulatory | 0.6 |
| MA-1_10 | MA-2_33 | biological | biological | 0.7 |
| MA-1_12 | MA-2_6 | regulatory | regulatory | 0.44 |
| MA-1_15 | MA-2_30 | regulatory | regulatory | 0.57 |
| MA-1_16 | MA-2_35 | biological | biological | 0.65 |
| MA-1_18 | MA-2_19 | biological | biological | 0.45 |
| MA-1_27 | MA-2_25 | biological | biological | 0.5 |
| MA-1_34 | MA-2_15 | regulatory | regulatory | 0.5 |
| MA-1_39 | MA-2_46 | regulatory | regulatory | 0.53 |
| MA-1_49 | MA-2_0 | regulatory | regulatory | 0.25 |
| MA-1_52 | MA-2_4 | regulatory | regulatory | 0.42 |
| MA-1_60 | MA-2_42 | biological | uncharacterized | 0.49 |
| MA-1_62 | MA-2_52 | uncharacterized | uncharacterized | 0.58 |
| MA-1_65 | MA-2_1 | biological | uncharacterized | 0.69 |
| MA-1_68 | MA-2_10 | regulatory | regulatory | 0.37 |
| MA-1_71 | MA-2_27 | uncharacterized | uncharacterized | 0.52 |
| MA-1_72 | MA-2_2 | regulatory | regulatory | 0.66 |
| MA-1_74 | MA-2_3 | uncharacterized | uncharacterized | 0.57 |
| MA-1_89 | MA-2_13 | regulatory | regulatory | 0.53 |
| MA-1_92 | MA-2_24 | uncharacterized | uncharacterized | 0.43 |
| MA-1_1 | MA-3_12 | regulatory | biological | 0.67 |
| MA-1_5 | MA-3_11 | regulatory | regulatory | 0.41 |
| MA-1_12 | MA-3_6 | regulatory | regulatory | 0.6 |

| | | | | |
|---|---|---|---|---|
| MA-1_15 | MA-3_15 | regulatory | regulatory | 0.54 |
| MA-1_34 | MA-3_1 | regulatory | regulatory | 0.63 |
| MA-1_49 | MA-3_9 | regulatory | regulatory | 0.21 |
| MA-1_67 | MA-3_21 | regulatory | regulatory | 0.68 |
| MA-1_68 | MA-3_27 | regulatory | regulatory | 0.56 |
| MA-1_72 | MA-3_30 | regulatory | regulatory | 0.7 |
| MA-1_82 | MA-3_17 | uncharacterized | uncharacterized | 0.69 |
| MA-2_0 | RNAseq-1_65 | regulatory | regulatory | 0.37 |
| MA-2_2 | RNAseq-1_22 | regulatory | regulatory | 0.66 |
| MA-2_4 | RNAseq-1_10 | regulatory | regulatory | 0.44 |
| MA-2_6 | RNAseq-1_46 | regulatory | regulatory | 0.38 |
| MA-2_7 | RNAseq-1_37 | regulatory | regulatory | 0.41 |
| MA-2_10 | RNAseq-1_33 | regulatory | regulatory | 0.32 |
| MA-2_12 | RNAseq-1_73 | regulatory | regulatory | 0.59 |
| MA-2_13 | RNAseq-1_4 | regulatory | regulatory | 0.43 |
| MA-2_15 | RNAseq-1_16 | regulatory | regulatory | 0.22 |
| MA-2_16 | RNAseq-1_74 | regulatory | regulatory | 0.51 |
| MA-2_17 | RNAseq-1_63 | regulatory | regulatory | 0.24 |
| MA-2_18 | RNAseq-1_17 | regulatory | regulatory | 0.43 |
| MA-2_20 | RNAseq-1_8 | regulatory | regulatory | 0.64 |
| MA-2_22 | RNAseq-1_36 | uncharacterized | genomic | 0.57 |
| MA-2_23 | RNAseq-1_1 | uncharacterized | biological | 0.54 |
| MA-2_25 | RNAseq-1_80 | biological | biological | 0.68 |
| MA-2_28 | RNAseq-1_0 | regulatory | regulatory | 0.51 |
| MA-2_29 | RNAseq-1_57 | regulatory | regulatory | 0.64 |
| MA-2_30 | RNAseq-1_58 | regulatory | regulatory | 0.56 |
| MA-2_31 | RNAseq-1_75 | regulatory | regulatory | 0.53 |
| MA-2_33 | RNAseq-1_25 | biological | regulatory | 0.62 |
| MA-2_36 | RNAseq-1_3 | uncharacterized | biological | 0.65 |
| MA-2_37 | RNAseq-1_55 | regulatory | regulatory | 0.6 |
| MA-2_41 | RNAseq-1_64 | regulatory | regulatory | 0.33 |
| MA-2_45 | RNAseq-1_87 | regulatory | regulatory | 0.62 |
| MA-2_46 | RNAseq-1_41 | regulatory | regulatory | 0.63 |
| MA-2_50 | RNAseq-1_9 | regulatory | regulatory | 0.69 |
| MA-2_0 | RNAseq-2_24 | regulatory | regulatory | 0.5 |
| MA-2_4 | RNAseq-2_34 | regulatory | regulatory | 0.52 |
| MA-2_6 | RNAseq-2_25 | regulatory | regulatory | 0.3 |
| MA-2_7 | RNAseq-2_47 | regulatory | regulatory | 0.58 |
| MA-2_8 | RNAseq-2_32 | uncharacterized | regulatory | 0.55 |
| MA-2_10 | RNAseq-2_39 | regulatory | regulatory | 0.41 |
| MA-2_13 | RNAseq-2_37 | regulatory | regulatory | 0.5 |

| | | | | |
|---|---|---|---|---|
| MA-2_14 | RNAseq-2_14 | regulatory | regulatory | 0.56 |
| MA-2_15 | RNAseq-2_45 | regulatory | regulatory | 0.52 |
| MA-2_17 | RNAseq-2_6 | regulatory | regulatory | 0.48 |
| MA-2_18 | RNAseq-2_5 | regulatory | regulatory | 0.6 |
| MA-2_20 | RNAseq-2_31 | regulatory | regulatory | 0.62 |
| MA-2_28 | RNAseq-2_35 | regulatory | regulatory | 0.67 |
| MA-2_31 | RNAseq-2_4 | regulatory | uncharacterized | 0.61 |
| MA-2_41 | RNAseq-2_44 | regulatory | regulatory | 0.32 |
| MA-2_0 | MA-3_9 | regulatory | regulatory | 0.23 |
| MA-2_6 | MA-3_6 | regulatory | regulatory | 0.62 |
| MA-2_7 | MA-3_10 | regulatory | regulatory | 0.47 |
| MA-2_10 | MA-3_27 | regulatory | regulatory | 0.47 |
| MA-2_14 | MA-3_19 | regulatory | regulatory | 0.6 |
| MA-2_15 | MA-3_1 | regulatory | regulatory | 0.44 |
| MA-2_28 | MA-3_28 | regulatory | regulatory | 0.58 |
| MA-2_30 | MA-3_15 | regulatory | regulatory | 0.62 |
| MA-2_31 | MA-3_30 | regulatory | regulatory | 0.5 |
| MA-2_41 | MA-3_11 | regulatory | regulatory | 0.38 |
| MA-2_43 | MA-3_24 | uncharacterized | uncharacterized | 0.63 |
| MA-2_51 | MA-3_29 | regulatory | regulatory | 0.69 |
| MA-3_1 | RNAseq-1_16 | regulatory | regulatory | 0.43 |
| MA-3_2 | RNAseq-1_73 | regulatory | regulatory | 0.68 |
| MA-3_6 | RNAseq-1_46 | regulatory | regulatory | 0.57 |
| MA-3_9 | RNAseq-1_65 | regulatory | regulatory | 0.31 |
| MA-3_10 | RNAseq-1_37 | regulatory | regulatory | 0.58 |
| MA-3_11 | RNAseq-1_64 | regulatory | regulatory | 0.37 |
| MA-3_12 | RNAseq-1_82 | biological | regulatory | 0.68 |
| MA-3_15 | RNAseq-1_58 | regulatory | regulatory | 0.48 |
| MA-3_17 | RNAseq-1_79 | uncharacterized | uncharacterized | 0.57 |
| MA-3_21 | RNAseq-1_27 | regulatory | regulatory | 0.6 |
| MA-3_23 | RNAseq-1_11 | uncharacterized | regulatory | 0.68 |
| MA-3_25 | RNAseq-1_24 | uncharacterized | regulatory | 0.69 |
| MA-3_27 | RNAseq-1_33 | regulatory | regulatory | 0.54 |
| MA-3_28 | RNAseq-1_0 | regulatory | regulatory | 0.26 |
| MA-3_30 | RNAseq-1_22 | regulatory | regulatory | 0.62 |
| MA-3_31 | RNAseq-1_59 | regulatory | regulatory | 0.61 |
| MA-3_1 | RNAseq-2_45 | regulatory | regulatory | 0.61 |
| MA-3_6 | RNAseq-2_25 | regulatory | regulatory | 0.55 |
| MA-3_9 | RNAseq-2_24 | regulatory | regulatory | 0.46 |
| MA-3_10 | RNAseq-2_47 | regulatory | regulatory | 0.68 |
| MA-3_11 | RNAseq-2_44 | regulatory | regulatory | 0.35 |

| | | | | |
|---|---|---|---|---|
| MA-3_17 | RNAseq-2_8 | uncharacterized | uncharacterized | 0.57 |
| MA-3_19 | RNAseq-2_14 | regulatory | regulatory | 0.59 |
| MA-3_21 | RNAseq-2_0 | regulatory | regulatory | 0.57 |
| MA-3_27 | RNAseq-2_39 | regulatory | regulatory | 0.61 |
| MA-3_28 | RNAseq-2_35 | regulatory | regulatory | 0.54 |

**Supplementary Table 3:** Reciprocal best hits between i-modulons from the individual transcriptomics datasets and i-modulons from the combined compendium

| **I-modulon from Combined Compendium** | **I-modulon from Individual Dataset** | **Type 1** | **Type 2** | **Distance** |
|---|---|---|---|---|
| Compendium_0 | RNAseq-1_24 | regulatory | regulatory | 0.73 |
| Compendium_1 | MA-1_36 | regulatory | regulatory | 0.56 |
| Compendium_1 | MA-2_39 | regulatory | regulatory | 0.46 |
| Compendium_1 | RNAseq-2_50 | regulatory | regulatory | 0.51 |
| Compendium_2 | RNAseq-1_12 | genomic | genomic | 0.94 |
| Compendium_3 | MA-1_58 | regulatory | regulatory | 0.46 |
| Compendium_3 | MA-3_31 | regulatory | regulatory | 0.61 |
| Compendium_3 | RNAseq-1_59 | regulatory | regulatory | 0.88 |
| Compendium_3 | RNAseq-2_41 | regulatory | regulatory | 0.3 |
| Compendium_4 | RNAseq-1_34 | regulatory | regulatory | 0.95 |
| Compendium_5 | RNAseq-1_26 | regulatory | regulatory | 0.96 |
| Compendium_6 | RNAseq-1_31 | uncharacterized | uncharacterized | 0.84 |
| Compendium_9 | RNAseq-1_68 | genomic | genomic | 0.84 |
| Compendium_10 | MA-2_15 | regulatory | regulatory | 0.79 |
| Compendium_10 | MA-3_1 | regulatory | regulatory | 0.56 |
| Compendium_10 | RNAseq-1_16 | regulatory | regulatory | 0.92 |
| Compendium_11 | RNAseq-1_62 | uncharacterized | biological | 0.62 |
| Compendium_11 | RNAseq-2_42 | uncharacterized | regulatory | 0.33 |
| Compendium_12 | RNAseq-2_17 | regulatory | regulatory | 0.4 |
| Compendium_13 | MA-1_67 | uncharacterized | regulatory | 0.37 |
| Compendium_13 | MA-2_21 | uncharacterized | uncharacterized | 0.68 |
| Compendium_14 | RNAseq-1_47 | regulatory | regulatory | 0.93 |
| Compendium_17 | MA-1_68 | regulatory | regulatory | 0.71 |
| Compendium_17 | MA-2_10 | regulatory | regulatory | 0.75 |
| Compendium_17 | MA-3_27 | regulatory | regulatory | 0.46 |
| Compendium_17 | RNAseq-1_33 | regulatory | regulatory | 0.9 |
| Compendium_17 | RNAseq-2_39 | regulatory | regulatory | 0.62 |
| Compendium_18 | RNAseq-1_5 | regulatory | regulatory | 0.91 |
| Compendium_19 | RNAseq-2_40 | regulatory | regulatory | 0.53 |

| | | | | |
|---|---|---|---|---|
| Compendium_20 | MA-2_45 | regulatory | regulatory | 0.42 |
| Compendium_20 | RNAseq-1_87 | regulatory | regulatory | 0.63 |
| Compendium_21 | MA-3_23 | uncharacterized | uncharacterized | 0.48 |
| Compendium_22 | RNAseq-1_28 | genomic | genomic | 0.89 |
| Compendium_23 | MA-1_17 | biological | genomic | 0.32 |
| Compendium_23 | MA-2_23 | biological | uncharacterized | 0.6 |
| Compendium_23 | RNAseq-1_1 | biological | biological | 0.84 |
| Compendium_24 | RNAseq-1_23 | regulatory | regulatory | 0.92 |
| Compendium_25 | MA-1_55 | regulatory | biological | 0.45 |
| Compendium_26 | MA-1_11 | regulatory | regulatory | 0.88 |
| Compendium_27 | MA-1_60 | uncharacterized | biological | 0.66 |
| Compendium_27 | MA-2_42 | uncharacterized | uncharacterized | 0.67 |
| Compendium_28 | MA-1_72 | regulatory | regulatory | 0.47 |
| Compendium_28 | MA-2_2 | regulatory | regulatory | 0.38 |
| Compendium_28 | RNAseq-1_22 | regulatory | regulatory | 0.64 |
| Compendium_28 | RNAseq-2_7 | regulatory | regulatory | 0.54 |
| Compendium_29 | RNAseq-1_56 | regulatory | regulatory | 0.95 |
| Compendium_30 | MA-2_18 | regulatory | regulatory | 0.7 |
| Compendium_30 | RNAseq-1_17 | regulatory | regulatory | 0.85 |
| Compendium_30 | RNAseq-2_5 | regulatory | regulatory | 0.65 |
| Compendium_32 | RNAseq-2_29 | genomic | genomic | 0.71 |
| Compendium_33 | RNAseq-1_40 | regulatory | regulatory | 0.93 |
| Compendium_34 | MA-1_38 | regulatory | regulatory | 0.43 |
| Compendium_34 | RNAseq-1_37 | regulatory | regulatory | 0.62 |
| Compendium_34 | RNAseq-2_47 | regulatory | regulatory | 0.52 |
| Compendium_35 | MA-2_8 | uncharacterized | uncharacterized | 0.74 |
| Compendium_35 | RNAseq-2_32 | uncharacterized | regulatory | 0.52 |
| Compendium_36 | MA-1_13 | biological | biological | 0.46 |
| Compendium_36 | MA-3_29 | biological | regulatory | 0.33 |
| Compendium_37 | RNAseq-1_30 | regulatory | regulatory | 0.61 |
| Compendium_40 | MA-1_49 | regulatory | regulatory | 0.67 |
| Compendium_40 | MA-2_0 | regulatory | regulatory | 0.66 |
| Compendium_40 | MA-3_9 | regulatory | regulatory | 0.65 |
| Compendium_40 | RNAseq-1_65 | regulatory | regulatory | 0.79 |

| | | | | |
|---|---|---|---|---|
| Compendium_40 | RNAseq-2_24 | regulatory | regulatory | 0.52 |
| Compendium_42 | MA-2_50 | regulatory | regulatory | 0.53 |
| Compendium_42 | RNAseq-1_9 | regulatory | regulatory | 0.69 |
| Compendium_43 | MA-1_93 | uncharacterized | genomic | 0.66 |
| Compendium_44 | MA-1_78 | uncharacterized | uncharacterized | 0.52 |
| Compendium_44 | MA-2_19 | uncharacterized | biological | 0.6 |
| Compendium_45 | MA-2_49 | uncharacterized | uncharacterized | 0.47 |
| Compendium_45 | RNAseq-2_33 | uncharacterized | uncharacterized | 0.7 |
| Compendium_46 | RNAseq-1_15 | genomic | genomic | 0.9 |
| Compendium_47 | RNAseq-1_29 | regulatory | regulatory | 0.93 |
| Compendium_48 | MA-1_43 | regulatory | uncharacterized | 0.31 |
| Compendium_48 | RNAseq-1_75 | regulatory | regulatory | 0.8 |
| Compendium_48 | RNAseq-2_4 | regulatory | uncharacterized | 0.38 |
| Compendium_49 | RNAseq-1_77 | regulatory | regulatory | 0.67 |
| Compendium_49 | RNAseq-2_12 | regulatory | regulatory | 0.77 |
| Compendium_51 | RNAseq-1_90 | genomic | genomic | 0.74 |
| Compendium_52 | MA-1_16 | biological | biological | 0.55 |
| Compendium_52 | MA-2_35 | biological | biological | 0.57 |
| Compendium_53 | RNAseq-1_50 | genomic | genomic | 0.93 |
| Compendium_54 | MA-1_3 | regulatory | regulatory | 0.56 |
| Compendium_55 | MA-1_27 | regulatory | biological | 0.39 |
| Compendium_55 | MA-2_25 | regulatory | biological | 0.48 |
| Compendium_56 | RNAseq-1_53 | regulatory | regulatory | 0.74 |
| Compendium_56 | RNAseq-2_10 | regulatory | regulatory | 0.38 |
| Compendium_57 | MA-3_30 | regulatory | regulatory | 0.51 |
| Compendium_58 | RNAseq-1_67 | regulatory | regulatory | 0.9 |
| Compendium_59 | MA-2_22 | genomic | uncharacterized | 0.7 |
| Compendium_59 | RNAseq-1_36 | genomic | genomic | 0.76 |
| Compendium_60 | RNAseq-1_13 | regulatory | regulatory | 0.74 |
| Compendium_61 | MA-1_1 | regulatory | regulatory | 0.64 |
| Compendium_61 | MA-2_31 | regulatory | regulatory | 0.4 |
| Compendium_61 | MA-3_12 | regulatory | biological | 0.45 |
| Compendium_61 | RNAseq-1_82 | regulatory | regulatory | 0.5 |
| Compendium_63 | MA-1_34 | regulatory | regulatory | 0.47 |

| | | | | |
|---|---|---|---|---|
| Compendium_63 | MA-2_16 | regulatory | regulatory | 0.76 |
| Compendium_63 | RNAseq-1_74 | regulatory | regulatory | 0.58 |
| Compendium_63 | RNAseq-2_45 | regulatory | regulatory | 0.52 |
| Compendium_64 | RNAseq-1_38 | biological | biological | 0.84 |
| Compendium_65 | MA-1_70 | regulatory | regulatory | 0.6 |
| Compendium_66 | MA-1_52 | regulatory | regulatory | 0.79 |
| Compendium_66 | MA-2_4 | regulatory | regulatory | 0.73 |
| Compendium_66 | RNAseq-1_10 | regulatory | regulatory | 0.82 |
| Compendium_66 | RNAseq-2_34 | regulatory | regulatory | 0.66 |
| Compendium_69 | RNAseq-1_11 | regulatory | regulatory | 0.94 |
| Compendium_70 | MA-1_41 | biological | genomic | 0.6 |
| Compendium_71 | MA-1_74 | uncharacterized | uncharacterized | 0.52 |
| Compendium_71 | MA-2_3 | uncharacterized | uncharacterized | 0.57 |
| Compendium_73 | RNAseq-2_36 | regulatory | regulatory | 0.79 |
| Compendium_76 | RNAseq-2_2 | regulatory | regulatory | 0.85 |
| Compendium_77 | MA-1_56 | uncharacterized | uncharacterized | 0.52 |
| Compendium_78 | RNAseq-1_83 | regulatory | regulatory | 0.59 |
| Compendium_78 | RNAseq-2_26 | regulatory | regulatory | 0.84 |
| Compendium_79 | RNAseq-1_20 | regulatory | regulatory | 0.8 |
| Compendium_80 | RNAseq-1_43 | regulatory | regulatory | 0.83 |
| Compendium_81 | MA-2_38 | regulatory | regulatory | 0.46 |
| Compendium_81 | MA-3_10 | regulatory | regulatory | 0.46 |
| Compendium_82 | MA-2_36 | uncharacterized | uncharacterized | 0.51 |
| Compendium_82 | RNAseq-1_3 | uncharacterized | biological | 0.75 |
| Compendium_82 | RNAseq-2_51 | uncharacterized | uncharacterized | 0.43 |
| Compendium_83 | RNAseq-1_48 | regulatory | regulatory | 0.73 |
| Compendium_84 | MA-2_51 | regulatory | regulatory | 0.55 |
| Compendium_84 | RNAseq-2_23 | regulatory | regulatory | 0.39 |
| Compendium_85 | RNAseq-1_52 | genomic | genomic | 0.72 |
| Compendium_86 | MA-1_91 | biological | biological | 0.87 |
| Compendium_87 | MA-2_29 | regulatory | regulatory | 0.47 |
| Compendium_87 | RNAseq-1_57 | regulatory | regulatory | 0.85 |
| Compendium_87 | RNAseq-2_9 | regulatory | regulatory | 0.35 |
| Compendium_88 | MA-1_81 | genomic | biological | 0.38 |

| | | | | |
|---|---|---|---|---|
| Compendium_89 | MA-1_5 | regulatory | regulatory | 0.72 |
| Compendium_89 | MA-2_41 | regulatory | regulatory | 0.84 |
| Compendium_89 | MA-3_11 | regulatory | regulatory | 0.52 |
| Compendium_89 | RNAseq-1_64 | regulatory | regulatory | 0.68 |
| Compendium_89 | RNAseq-2_44 | regulatory | regulatory | 0.62 |
| Compendium_90 | RNAseq-1_7 | uncharacterized | uncharacterized | 0.84 |
| Compendium_91 | MA-2_17 | regulatory | regulatory | 0.83 |
| Compendium_91 | RNAseq-1_63 | regulatory | regulatory | 0.94 |
| Compendium_91 | RNAseq-2_6 | regulatory | regulatory | 0.66 |
| Compendium_93 | RNAseq-1_66 | regulatory | regulatory | 0.9 |
| Compendium_93 | RNAseq-2_18 | regulatory | regulatory | 0.84 |
| Compendium_94 | MA-1_10 | regulatory | biological | 0.51 |
| Compendium_94 | MA-2_33 | regulatory | biological | 0.44 |
| Compendium_94 | RNAseq-2_21 | regulatory | biological | 0.62 |
| Compendium_95 | MA-1_28 | biological | biological | 0.85 |
| Compendium_97 | RNAseq-1_71 | regulatory | regulatory | 0.89 |
| Compendium_99 | MA-1_59 | regulatory | uncharacterized | 0.44 |
| Compendium_99 | MA-2_24 | regulatory | uncharacterized | 0.34 |
| Compendium_100 | RNAseq-2_38 | biological | biological | 0.66 |
| Compendium_101 | MA-3_4 | regulatory | biological | 0.57 |
| Compendium_102 | MA-1_75 | regulatory | regulatory | 0.51 |
| Compendium_105 | MA-1_15 | regulatory | regulatory | 0.75 |
| Compendium_105 | MA-2_30 | regulatory | regulatory | 0.51 |
| Compendium_105 | MA-3_15 | regulatory | regulatory | 0.6 |
| Compendium_105 | RNAseq-1_58 | regulatory | regulatory | 0.93 |
| Compendium_106 | RNAseq-1_72 | regulatory | regulatory | 0.79 |
| Compendium_106 | RNAseq-2_3 | regulatory | regulatory | 0.48 |
| Compendium_107 | RNAseq-1_80 | uncharacterized | biological | 0.58 |
| Compendium_108 | MA-1_88 | regulatory | regulatory | 0.85 |
| Compendium_109 | MA-2_28 | regulatory | regulatory | 0.56 |
| Compendium_109 | MA-3_28 | regulatory | regulatory | 0.77 |
| Compendium_109 | RNAseq-1_0 | regulatory | regulatory | 0.96 |
| Compendium_109 | RNAseq-2_35 | regulatory | regulatory | 0.59 |
| Compendium_111 | RNAseq-1_25 | biological | regulatory | 0.36 |

| | | | | |
|---|---|---|---|---|
| Compendium_112 | MA-1_39 | regulatory | regulatory | 0.84 |
| Compendium_112 | MA-2_46 | regulatory | regulatory | 0.57 |
| Compendium_112 | RNAseq-1_41 | regulatory | regulatory | 0.72 |
| Compendium_112 | RNAseq-2_30 | regulatory | regulatory | 0.56 |
| Compendium_113 | RNAseq-2_16 | biological | genomic | 0.76 |
| Compendium_116 | MA-1_6 | regulatory | regulatory | 0.61 |
| Compendium_116 | MA-2_20 | regulatory | regulatory | 0.55 |
| Compendium_116 | RNAseq-1_8 | regulatory | regulatory | 0.63 |
| Compendium_116 | RNAseq-2_31 | regulatory | regulatory | 0.74 |
| Compendium_117 | MA-1_14 | biological | biological | 0.55 |
| Compendium_117 | RNAseq-1_35 | biological | biological | 0.6 |
| Compendium_118 | MA-1_85 | genomic | genomic | 0.73 |
| Compendium_119 | RNAseq-1_19 | regulatory | regulatory | 0.66 |
| Compendium_120 | MA-1_12 | regulatory | regulatory | 0.73 |
| Compendium_120 | MA-2_6 | regulatory | regulatory | 0.75 |
| Compendium_120 | MA-3_6 | regulatory | regulatory | 0.5 |
| Compendium_120 | RNAseq-1_46 | regulatory | regulatory | 0.84 |
| Compendium_120 | RNAseq-2_25 | regulatory | regulatory | 0.92 |
| Compendium_121 | MA-1_62 | uncharacterized | uncharacterized | 0.66 |
| Compendium_121 | MA-2_52 | uncharacterized | uncharacterized | 0.53 |
| Compendium_122 | MA-1_0 | regulatory | biological | 0.55 |
| Compendium_122 | MA-2_7 | regulatory | regulatory | 0.54 |
| Compendium_123 | MA-1_89 | regulatory | regulatory | 0.51 |
| Compendium_123 | MA-2_13 | regulatory | regulatory | 0.69 |
| Compendium_123 | RNAseq-1_4 | regulatory | regulatory | 0.86 |
| Compendium_123 | RNAseq-2_37 | regulatory | regulatory | 0.58 |
| Compendium_125 | RNAseq-1_89 | regulatory | regulatory | 0.72 |
| Compendium_126 | MA-1_102 | regulatory | regulatory | 0.69 |
| Compendium_127 | MA-1_32 | regulatory | regulatory | 0.36 |
| Compendium_127 | RNAseq-1_6 | regulatory | regulatory | 0.83 |
| Compendium_128 | MA-1_80 | regulatory | regulatory | 0.81 |
| Compendium_128 | RNAseq-1_76 | regulatory | regulatory | 0.65 |
| Compendium_130 | RNAseq-1_70 | regulatory | regulatory | 0.91 |
| Compendium_131 | RNAseq-1_45 | regulatory | regulatory | 0.92 |

| | | | | |
|---|---|---|---|---|
| Compendium_134 | RNAseq-1_18 | regulatory | regulatory | 0.89 |
| Compendium_136 | RNAseq-1_69 | genomic | genomic | 0.89 |
| Compendium_137 | RNAseq-1_51 | biological | biological | 0.8 |
| Compendium_138 | RNAseq-1_42 | genomic | genomic | 0.87 |
| Compendium_139 | RNAseq-1_2 | uncharacterized | uncharacterized | 0.82 |
| Compendium_140 | RNAseq-1_54 | regulatory | regulatory | 0.91 |
| Compendium_142 | RNAseq-1_60 | regulatory | regulatory | 0.8 |
| Compendium_143 | MA-2_48 | regulatory | regulatory | 0.49 |
| Compendium_144 | RNAseq-1_61 | regulatory | regulatory | 0.82 |
| Compendium_146 | RNAseq-1_85 | regulatory | regulatory | 0.82 |
| Compendium_147 | MA-1_82 | uncharacterized | uncharacterized | 0.45 |
| Compendium_147 | MA-3_17 | uncharacterized | uncharacterized | 0.4 |
| Compendium_147 | RNAseq-1_79 | uncharacterized | uncharacterized | 0.68 |
| Compendium_147 | RNAseq-2_8 | uncharacterized | uncharacterized | 0.71 |
| Compendium_148 | MA-1_21 | regulatory | regulatory | 0.34 |
| Compendium_148 | MA-2_37 | regulatory | regulatory | 0.57 |
| Compendium_148 | RNAseq-1_55 | regulatory | regulatory | 0.79 |
| Compendium_149 | MA-2_14 | regulatory | regulatory | 0.6 |
| Compendium_149 | MA-3_19 | regulatory | regulatory | 0.66 |
| Compendium_149 | RNAseq-2_14 | regulatory | regulatory | 0.73 |
| Compendium_150 | MA-1_57 | regulatory | biological | 0.38 |
| Compendium_151 | MA-2_32 | regulatory | regulatory | 0.66 |
| Compendium_152 | MA-2_12 | regulatory | regulatory | 0.54 |
| Compendium_152 | MA-3_2 | regulatory | regulatory | 0.41 |
| Compendium_152 | RNAseq-1_73 | regulatory | regulatory | 0.87 |
| Compendium_152 | RNAseq-2_43 | regulatory | regulatory | 0.38 |
| Compendium_153 | MA-1_37 | regulatory | uncharacterized | 0.39 |
| Compendium_153 | RNAseq-1_32 | regulatory | regulatory | 0.74 |
| Compendium_154 | MA-1_25 | regulatory | uncharacterized | 0.48 |
| Compendium_155 | RNAseq-1_44 | regulatory | regulatory | 0.92 |
| Compendium_156 | MA-1_35 | regulatory | regulatory | 0.49 |
| Compendium_157 | MA-3_21 | uncharacterized | regulatory | 0.3 |
| Compendium_157 | RNAseq-1_27 | uncharacterized | regulatory | 0.58 |
| Compendium_157 | RNAseq-2_0 | uncharacterized | regulatory | 0.54 |

| | | | | |
|---|---|---|---|---|
| Compendium_158 | RNAseq-1_14 | regulatory | regulatory | 0.84 |
| Compendium_158 | RNAseq-2_27 | regulatory | regulatory | 0.76 |
| Compendium_164 | MA-2_53 | biological | uncharacterized | 0.34 |
| Compendium_168 | MA-3_5 | regulatory | uncharacterized | 0.5 |
| Compendium_168 | RNAseq-1_81 | regulatory | regulatory | 0.57 |
| Compendium_171 | RNAseq-1_84 | biological | uncharacterized | 0.58 |
| Compendium_173 | RNAseq-2_49 | biological | uncharacterized | 0.59 |
| Compendium_174 | RNAseq-1_39 | biological | biological | 0.91 |
| Compendium_175 | RNAseq-1_86 | genomic | genomic | 0.41 |
| Compendium_177 | RNAseq-2_11 | uncharacterized | regulatory | 0.51 |
| Compendium_180 | MA-2_9 | uncharacterized | uncharacterized | 0.36 |

**Supplementary Table 4:** Reciprocal best hits between the two proteomics datasets and the RNAseq-1 dataset. Distances above 0.7 have been pruned.

| I-modulon 1 | I-modulon 2 | Type 1 | Type 2 | Distance |
|---|---|---|---|---|
| RNAseq-1_3 | Proteomics-1_1 | biological | regulatory | 0.61 |
| RNAseq-1_11 | Proteomics-1_10 | regulatory | uncharacterized | 0.65 |
| RNAseq-1_16 | Proteomics-1_0 | regulatory | uncharacterized | 0.58 |
| RNAseq-1_18 | Proteomics-1_5 | regulatory | regulatory | 0.65 |
| RNAseq-1_24 | Proteomics-1_2 | regulatory | uncharacterized | 0.66 |
| RNAseq-1_26 | Proteomics-1_11 | regulatory | regulatory | 0.61 |
| RNAseq-1_27 | Proteomics-1_6 | regulatory | regulatory | 0.67 |
| RNAseq-1_33 | Proteomics-1_13 | regulatory | regulatory | 0.58 |
| RNAseq-1_56 | Proteomics-1_8 | regulatory | regulatory | 0.42 |
| RNAseq-1_58 | Proteomics-1_17 | regulatory | regulatory | 0.43 |
| RNAseq-1_2 | Proteomics-2_16 | uncharacterized | uncharacterized | 0.68 |
| RNAseq-1_10 | Proteomics-2_12 | regulatory | biological | 0.69 |
| RNAseq-1_16 | Proteomics-2_1 | regulatory | regulatory | 0.58 |
| RNAseq-1_26 | Proteomics-2_11 | regulatory | regulatory | 0.37 |
| RNAseq-1_33 | Proteomics-2_4 | regulatory | regulatory | 0.66 |
| RNAseq-1_47 | Proteomics-2_6 | regulatory | uncharacterized | 0.6 |
| RNAseq-1_75 | Proteomics-2_9 | regulatory | regulatory | 0.64 |
| RNAseq-1_76 | Proteomics-2_3 | regulatory | regulatory | 0.54 |
| Proteomics-1_6 | Proteomics-2_3 | regulatory | regulatory | 0.36 |
| Proteomics-1_11 | Proteomics-2_11 | regulatory | regulatory | 0.6 |

**Supplementary Table 5:** Linear regression coefficients for the 10 RNAseq-1 ICs that loosely approximate a Proteomics-1 IC (See Figure5i)

| **I-modulon** | **Description** | **Linear Regression Coefficient** |
|---|---|---:|
| RNAseq-1_24 | Lrp | 0.38 |
| RNAseq-1_27 | Crp | 0.19 |
| RNAseq-1_58 | GlpR | 0.13 |
| RNAseq-1_83 | TrpR | -0.15 |
| RNAseq-1_63 | CusR | -0.15 |
| RNAseq-1_17 | ArgR | -0.17 |
| RNAseq-1_77 | TyrR | -0.18 |
| RNAseq-1_73 | MetJ | -0.19 |
| RNAseq-1_87 | leu-tRNA-mediated attenuation | -0.23 |
| RNAseq-1_4 | UTP-mediated attenuation | -0.23 |
